# Cross-family learning transfers thermal adaptation between enzymes

**DOI:** 10.64898/2026.10.26.761089

**Authors:** Sandra Viknander, Nikolaos Tatarakis, Xiaozhi Fu, Alexander Diaciuc, Filip Buric, Aleksej Zelezniak

## Abstract

Thermal adaptation has evolved independently across diverse protein families, yet whether the information underlying this adaptation can be transferred between evolutionarily unrelated proteins remains unknown. Here, we demonstrate that thermostability can be learned from divergent protein families and transferred to enzymes without relying on their evolutionary ancestry. We developed THOR (THermally Optimised Representations), a generative framework that learns background-specific sequence transformations between unpaired mesophilic and thermophilic sequence populations while preserving the identity of the starting protein. Across evolutionarily divergent enzyme families, THOR transferred thermostability without exposure to thermophilic homologues during training. In malate dehydrogenase, excluded from cross-family training, zero-shot designs increased apparent melting temperature by up to 15 °C, with gains reaching 36 °C following limited family-specific fine-tuning. Majority of designed variants retained activity following prolonged thermal challenge, while selected variants exhibited nearly threefold higher catalytic turnover than wild type. Molecular dynamics simulations revealed that comparable thermal phenotypes emerge through distinct, background-dependent changes in protein flexibility and stabilising interactions. Together, these findings demonstrate that thermal adaptation encodes transferable information beyond lineage-specific evolutionary solutions, revealing evolution as a source of reusable adaptive principles for engineering phenotypes in proteins whose evolutionary lineages have never acquired them.

## Introduction

Enzymes must remain folded and functional within the environmental conditions in which they operate, making temperature one of the major selective pressures shaping protein evolution. Thermophilic organisms demonstrate that essentially the same biochemical functions can be maintained at temperatures that destabilise many mesophilic proteins, and these naturally evolved solutions have long provided principles for engineering more robust biocatalysts^1,2^. Yet natural evolution does not encode thermal adaptation in isolation, i.e. the sequence of a thermophilic enzyme reflects not only adaptation to temperature, but also phylogenetic history and other environmental, genomic and functional constraints acting during its evolution ^3,4^. Consequently, sequence differences between mesophilic and thermophilic homologues can conflate temperature-associated adaptation with lineage-specific evolutionary variation.

In theory, repeated independent transitions between mesophilic and thermophilic states within the same protein family could help uncover specific temperature-associated sequence changes from lineage-specific variation^4,5^. In practice, available sequences are strongly biased towards mesophilic organisms, leaving few highly divergent thermophilic lineages within most protein families for such comparisons^6^. Thermal adaptation is therefore repeatedly observed across the protein universe, but poorly represented within individual evolutionary lineages. Yet independent thermophilic lineages exhibit recurring amino-acid and substitutional biases, indicating that temperature adaptation can leave shared sequence signatures across distinct evolutionary backgrounds ^7,8^. Although individual substitutions and structural mechanisms may differ between lineages, repeated adaptation to the same environmental challenge may therefore encode transferable information that extends beyond the evolutionary background in which it arose.

Recent work supports the idea that evolutionary sequence variation contains separable components associated with different protein phenotypes. Large-scale measurements of protein stability have shown that natural amino-acid preferences cannot be explained by folding stability alone, consistent with proteins evolving under multiple simultaneous constraints^9^. Combinatorial sequence perturbations further indicate that substantial components of protein stability can arise from relatively simple mutational contributions rather than requiring dense networks of lineage-specific epistasis^10^. Consistent with this, Pinney et al. demonstrated that temperature adaptation can involve parallel molecular mechanisms across distinct evolutionary backgrounds, identifying temperature-associated residues across more than 1,000 bacterial enzyme families^11^. Recent experimental evidence further suggests that the sequence determinants of protein stability and function are partly separable: preferences for specific amino acids at individual positions primarily favour folding stability, whereas evolutionary correlations between residues play a greater role in maintaining catalytic activity^12^. Together, these observations suggest that components of stability-related sequence information can be distinguished from other evolutionary traces encoded in natural proteins. However, these studies largely infer such information within homologous protein families. Yet, whether an adaptive phenotype can be learned across unrelated protein families and transferred to enzymes with no evolutionary history of thermal adaptation remains unknown.

Most established protein-engineering strategies operate within, or are informed by, the evolutionary and structural context of the target protein. Directed evolution and deep mutational scanning explore its local mutational neighbourhood^13,14^; consensus design, ancestral reconstruction and other evolution-informed approaches exploit sequence variation among homologues^15,16^; and physics-based and structure-guided methods identify stabilising changes using the target fold or energetic model^17–19^. More recently, generative sequence models and inverse-folding approaches have greatly expanded the sequence space that can be explored and can generate highly stable proteins or improve the stability of existing enzymes^20–23^. These approaches can be remarkably effective, but they primarily identify stabilising solutions using evolutionary, structural or property information available for the target protein or its surrounding sequence space. They do not ask whether adaptive information can be isolated across unrelated evolutionary backgrounds and transferred into a protein family in which that phenotype was never observed. More broadly, resolving this question would establish whether evolution can be mined not only for individual solutions, but for reusable principles of adaptation.

Here, we asked whether thermal adaptation can be learned independently of the evolutionary history of the target protein family. We developed THOR (THermally Optimised Representations), a generative framework trained on ∼17 million mesophilic and thermophilic protein sequences that learns transformations between unpaired sequence populations while preserving the identity of the starting protein. Thermophilic homologues from held-out target families were excluded from training, forcing THOR to learn thermal-adaptation information from unrelated evolutionary backgrounds rather than matched mesophile–thermophile homologues or family-specific thermostability measurements. The resulting framework tests whether thermal-adaptation information can generalise to protein families in which the thermophilic phenotype was not represented during training. Across protein families excluded from training, THOR-generated sequences acquired thermophile-associated amino-acid usage and residue-conditioned substitution patterns while retaining their family-specific sequence context. We tested whether this information could confer thermostability on an unseen enzyme family using three evolutionarily divergent malate dehydrogenase (MDH) lineages, that were entirely excluded from training. Of the 44 designs from three lineages, 56% exhibited increases in apparent melting temperature of at least 5 °C, with gains reaching 37 °C, while 74% retained enzymatic activity following prolonged thermal exposure at temperatures exceeding the melting temperature of their corresponding wild-type enzymes. Selected variants also reached turnover numbers (*k_cat_*) up to threefold higher than wild type, demonstrating that increased thermal robustness remained compatible with high catalytic turnover. Molecular dynamics simulations further showed that, even within the same lineage, comparable thermal phenotypes emerged through distinct redistributions of protein dynamics rather than convergence on a single stabilisation mechanism. Together, these results show that thermal adaptation contains transferable sequence information that generalises across evolutionary lineages and can be introduced into new enzyme backgrounds without retracing the natural evolutionary route to thermostability.

## Results

### The enzyme universe is sparsely populated with thermostable catalysts

To assess how much natural sequence information is available for learning thermal adaptation within protein families, we examined two complementary sources of temperature information. Protein melting temperature (*T_m_*) provides a related measure of thermal stability but is experimentally available for relatively few proteins^24^. Organism growth temperature, which we refer to here as environmental temperature (ET), is available at substantially greater scale^6^ and correlates strongly with experimentally measured protein *T*_m_ ^11^(Pearson *r=0.94*; Supplementary Fig. 1), supporting its use as a broader indicator of thermal adaptation. We therefore combined experimental *T_m_* values from the Meltome Atlas^25^ with proteins annotated by ET. After redundancy filtering, the Meltome Atlas contained ∼48,000 proteins and the ET dataset ∼2.7 million sequences (Methods M1). Together, these represent less than 1% of UniProt and are strongly biased towards mesophilic conditions (Fig. 1A). Only ∼3,300 Meltome proteins have a *T_m_* above 60 °C, while ∼93,000 ET-annotated sequences originate from organisms growing above 45 °C (Fig. 1B).

**Figure 1.**
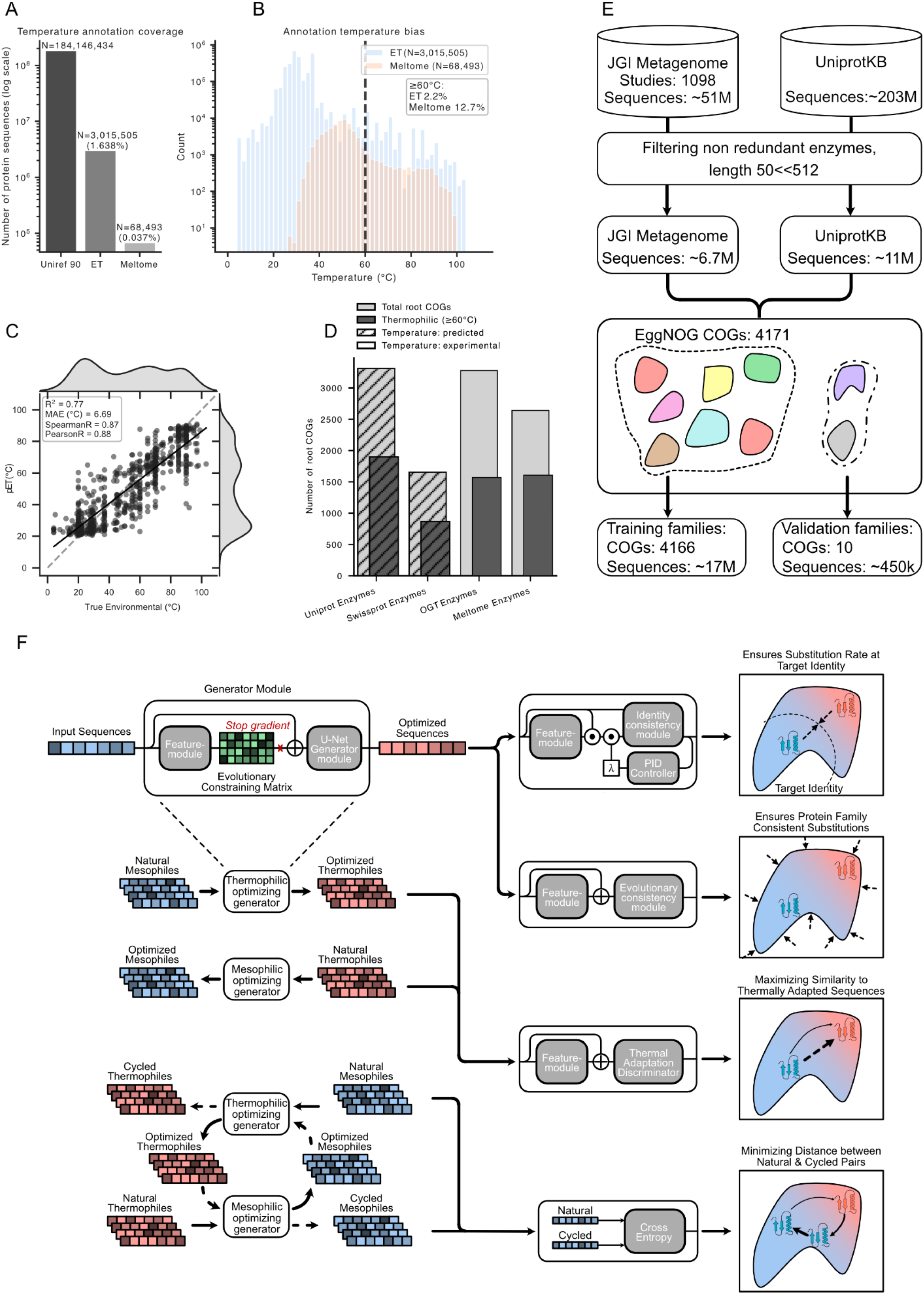
| Learning transferable thermal adaptation from sparse protein temperature data. **A**, Coverage of environmental temperatures (ET) annotations and experimentally measured protein melting temperatures across large protein sequence databases. **B**, Temperature distributions of ET-annotated enzymes and experimentally measured melting temperatures, highlighting the strong bias towards mesophilic conditions. **C**, Performance of the sequence-based temperature predictor on a uniformly sampled temperature test set, showing agreement between predicted and measured temperatures. **D**, Distribution of temperature information across EggNOG COG protein families, illustrating the limited representation of thermophilic enzymes within individual families. E, Construction of the training and held-out validation datasets from JGI metagenomes and UniProtKB. Protein sequences were filtered, assigned to EggNOG COGs and partitioned such that ten protein families were completely excluded from model training. **F**, THOR architecture for learning bidirectional transformations between mesophilic and thermophilic sequence states from unpaired protein sequences. The generators propose substitutions under evolutionary constraints, while identity consistency controls the extent of sequence change, evolutionary consistency favours substitutions compatible with the target protein family, and a thermal adaptation discriminator drives sequences towards the desired thermal state. Cycle consistency requires the original sequence to be recoverable after the forward and reverse transformations, thereby discouraging mode collapse and preserving sequence-specific information. COG-Clusters of Orthologous Groups; ET - Environmental Temperature.

Because experimentally measured *T_m_* and organism-level ET annotations cover only a small fraction of known protein sequences, we trained a sequence-based regression model to predict environmental temperature (pET) from amino-acid sequence (Methods M2). The model achieved an *R*^2^ =0.77 and a median absolute error of 6.69 °C on an independent test set (Fig. 1C, Supplementary Table 1), allowing us to estimate the distribution of high-temperature-associated sequences across the much larger UniProt database^26^. We then annotated UniProt with pET and examined their representation within enzyme families. Of 3311 clusters of orthologous groups (COGs) (Methods M1), only 57% contained at least a single sequence with pET above 60 °C (Fig. 1D), with a median of only 1 thermophilic sequence (Supplementary Fig. 2). Furthermore, substantial sequence diversity was observed within individual COGs, with a median pairwise sequence identity of only 26.1% between thermophilic and mesophilic sequences (Supplementary Fig. 3). Together, these findings highlight the sparse representation of thermophilic sequences across a highly diverse protein sequence space. Together, these data show that current thermophilic sequence diversity is sparsely represented within individual protein families.

### Learning thermal adaptation across unrelated evolutionary backgrounds

The sparse representation of thermophilic proteins within individual families limits the sequence information available for separating thermal adaptation from lineage-specific evolutionary variation. We therefore asked whether sequence information associated with thermal adaptation could instead be learned across unrelated evolutionary backgrounds. To address this, we developed THOR (THermally Optimised Representations), which treats thermal adaptation as a transformation between mesophilic and thermophilic sequence populations. Rather than identifying a particular thermophilic template or evolutionary trajectory, THOR learns sequence changes that distinguish these populations and applies them in the context of a target protein while preserving its identity.

To learn thermal adaptation across diverse evolutionary backgrounds, we assembled a dataset of mesophilic and thermophilic sequence populations. Because high-temperature proteins are poorly represented in curated databases (Fig.1D), we expanded the thermophilic population using 1,098 metagenomic samples collected from environments above 60 °C, comprising more than 50 million enzyme-coding genes across (Fig. 1E). After removal of signal peptides and clustering at 95% sequence identity, these data yielded 6.6 million putative thermophilic sequences, which were combined with 10.6 million mesophilic sequences from Uniprot enzymes (Methods M1). To test whether thermal adaptation could generalise beyond the families in which it was observed, we reserved ten protein families for evaluation (Supplementary Table 2) and removed their orthologues from both UniProt and the metagenomic training sets. The resulting ∼17-million-sequence dataset therefore contained extensive thermal-adaptation information across unrelated proteins while excluding the thermophilic phenotype from the families used for subsequent evaluation.

THOR learns this transformation using an unpaired, cycle-consistent generative framework inspired by domain translation methods^27^ (Fig. 1F, Supplementary Fig. 4-6). Mesophilic and thermophilic sequences are sampled independently during training, with homologous mesophile–thermophile pairs excluded, so the model is never provided with a direct evolutionary correspondence between the two populations. Two reciprocal generators map sequences between mesophilic and thermophilic populations, while discriminators encourage generated sequences to acquire characteristics of the target population (Methods M3). Cycle consistency requires sequences transformed into the opposite thermal domain and back to reconstruct their starting sequence, discouraging collapse onto generic thermophile-like solutions and preserving input-specific sequence information. An additional evolutionary loss derived from protein language-model probabilities penalises substitutions that are unlikely in the context of the starting protein^22,28^. The introduced proportional–integral–derivative (PID) controller dynamically adjusts this constraint to maintain a defined sequence identity to the starting protein (Methods M3). Together, these components force THOR to learn thermal-adaptation patterns shared across unrelated sequence backgrounds rather than transformations between homologous mesophile–thermophile pairs. To monitor whether THOR shifted generated sequences towards high-temperature-associated sequence space, we used the sequence-based pET model introduced above (Fig. 1C) as an independent readout. To note, pET model was not included in the THOR training objective and therefore provided no supervision for sequence generation and was just used for monitoring of training.

We next asked how much sequence change was required to produce a thermal shift while retaining the starting protein background. Across target identities of 70–95%, THOR consistently reached the specified sequence identity while increasing pET (Fig. 2A). Even at 95% identity, generated sequences showed average predicted increases exceeding 15 °C. The predicted thermal shift largely plateaued after ∼85% identity (Wilcoxon rank-sum test e.g. 80 vs 75%, p-value=0.43), with greater divergence producing little additional increase in pET (Supplementary Fig. 7). We therefore selected the 80% identity model for subsequent analyses and experimental validation. To assess training convergence, we additionally compared the model’s output probability distributions across training checkpoints using Jensen–Shannon divergence on a set of validation sequences (Supplementary Fig. 8). The increasing similarity between successive checkpoints, together with the stabilization of the target sequence identity (Fig. 2A).

**Figure 2.**
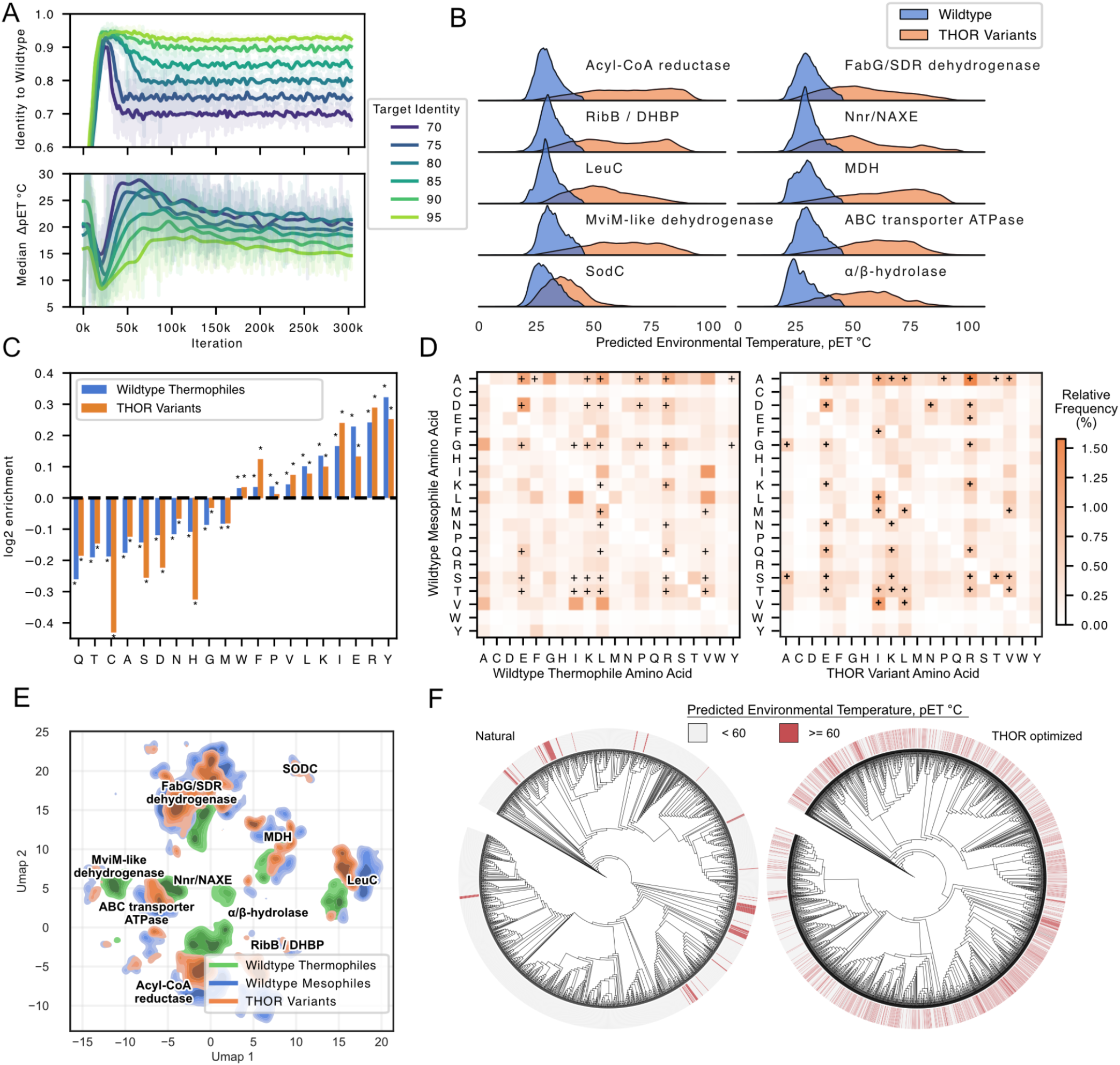
| THOR generalises thermal-adaptation patterns across unseen protein families. A, Training dynamics across prescribed target identities. Generated sequences converge towards the requested identity to the wild type while retaining increases in predicted environmental temperature, showing that sequence divergence can be constrained across different identity targets. **B**, Predicted environmental-temperature (pET) distributions for wild-type sequences and THOR variants across ten protein families excluded from training. THOR shifts sequences towards higher predicted temperatures across diverse protein families. **C**, Amino-acid enrichments in natural thermophiles and THOR variants relative to mesophilic wild types, revealing shared thermophile-associated compositional trends. * indicates BH adjusted p-value <0.01 using fisher’s exact test **D,** Substitution-frequency matrices comparing amino-acid changes observed between natural mesophilic and thermophilic proteins with those introduced by THOR. Crosses indicate enriched substitutions (BH adjusted p-value <0.01 using binomial test) for natural sequences (left panel) for THOR variants (right panel) + indicates 41 most frequent substitutions, highlighting overlap between naturally evolved and model-generated thermal adaptation patterns. **E,** UMAP projection of sequence representations for mesophilic wild types, thermophilic wild types and THOR variants across the held-out families. THOR variants remain within family-specific sequence neighbourhoods while extending towards thermophile-associated regions of sequence space. **F,** Phylogenetic distribution of wild-type proteins and THOR variants, coloured by predicted environmental temperature. Generated variants remain associated with their underlying sequence lineages while acquiring higher predicted thermal phenotypes, indicating that THOR transfers thermal adaptation without retracing natural thermophilic evolutionary trajectories (Supplementary Fig. 14 for other families).

### Thermal adaptation generalises across enzyme families

We first tested whether the learned transformation generalised zero-shot to protein families that were entirely excluded from training data (Fig. 1E). Across ten held-out orthogroups, THOR consistently shifted generated sequences towards higher predicted environmental temperatures, with a median increase in pET of 26 ± 8 °C (s.d.) relative to their mesophilic starting sequences (Fig. 2B). The generated sequences also showed significantly greater variation in predicted thermal shift than their corresponding wild types (Δs.d. = 10.6 °C, Wilcoxon rank-sum test, p-value = 0.00097), indicating that the magnitude of transfer depended on the sequence background rather than producing a uniform thermophilic shift. Because thermophilic homologues from these families were absent from training, these changes could not arise from reproducing family-specific thermophilic examples. Instead, the results indicate that thermal-adaptation information learned across unrelated evolutionary backgrounds can transfer zero-shot into previously unseen protein families.

We next asked whether the transferred sequences reproduced sequence features independently associated with natural thermophily. Across the held-out protein families, natural thermophilic sequences showed characteristic shifts in amino-acid usage relative to mesophilic enzymes, including enrichment of charged and hydrophobic residues and depletion of several polar and small residues (Fig. 2C). THOR-generated variants reproduced the direction of most of these changes despite having no thermophilic examples from the corresponding families during training. The strongest shifts included depletion of cysteine and histidine and enrichment of residues such as glutamate, lysine, leucine and arginine. Thus, cross-family transfer recapitulated known amino-acid usage biases associated with natural thermophily ^8,29^, rather than introducing arbitrary compositional changes.

Beyond these global compositional shifts, natural thermal adaptation also produces characteristic residue-specific substitution patterns. We therefore compared mesophile-to-thermophile substitutions in natural proteins with those introduced by THOR across the held-out families (Fig. 2D, Methods M4). Natural thermophilic sequences showed a sparse, non-random substitution profile, with enrichment of specific residue transitions rather than uniform replacement towards thermophile-enriched amino acids. THOR partially recapitulated this structure, including frequent substitutions such as A→K/L/R, D→E/R, G→E/R, Q→E/R and S/T→K/L/R. Across all residue pairs, the THOR and natural mesophile-to-thermophile substitution matrices were significantly correlated (average Pearson’s *r* = 0.80, p-value = 1.85e-85) (Supplementary Fig. 9) indicating shared substitutional biases. However, the matrices were far from identical, showing that THOR did not simply reproduce the natural mesophile-to-thermophile substitution programme. Consistent with this, THOR introduced distinct substitution profiles across divergent wild-type backgrounds from the same family (Supplementary Fig. 10,11), indicating that the transferred thermal signal was realised in a background-dependent manner rather than through a fixed set of thermophilic mutations. To assess whether these improvements could be attributed to the underlying ESM sequence probabilities or changes in amino-acid composition alone, we compared THOR-generated sequences against variants sampled from ESM and two random substitution controls (Supplementary Fig. 12). THOR consistently produced significantly higher predicted temperatures across protein families, supporting a contribution from the learned thermal adaptation model beyond these baseline sampling strategies.Together, these results show that the transferred signal extends beyond bulk amino-acid usage to residue-conditioned, background-dependent sequence changes that resemble natural thermal adaptation.

To examine how these residue-conditioned, background-dependent substitutions reshape sequence space, we embedded natural mesophilic and thermophilic proteins together with THOR variants using ESM representations and visualised them with UMAP^30^ (Fig. 2E, Methods M5). Within each protein family, THOR variants expanded beyond the mesophilic sequence population towards regions increasingly associated with natural thermophiles, while remaining distinct from the naturally evolved thermophilic sequences. Rather than converging on a common thermophilic solution, different families expanded along distinct regions of their local sequence landscapes. Thus, cross-family transfer opens thermophile-associated sequence space from multiple evolutionary starting points without reproducing the natural trajectories that occupy it.

We next examined how this expansion was distributed across the phylogeny. Natural thermophilic sequences were confined to a subset of clades, whereas THOR generated high-pET variants across a broader range of phylogenetic backgrounds (Fig. 2F, Supplementary Fig. 13-15, Methods M6). The number of clades containing at least one sequence with pET ≥60 °C increased significantly after THOR transformation (144 versus 413; , McNemar’s exact test, p-value < 1e-16). Moreover, high-pET THOR variants remained associated with their corresponding wild-type lineages rather than converging towards existing natural thermophilic clades. As a control for sequence divergence alone, ESM-sampled and random variants with the same average number of substitutions produced significantly weaker enrichment of high-pET sequences than THOR variants (3.6-fold lower for ESM-sampled; p-value = 0.0039). Thus, THOR introduces thermophile-associated properties locally across distinct evolutionary backgrounds rather than redirecting sequences towards existing thermophilic lineages.

### Thermostability can be transferred across evolutionarily divergent enzyme families

To test whether cross-family thermal-adaptation information could produce an experimentally measurable phenotype, we selected malate dehydrogenase (MDH) as an unseen enzyme family. MDH sequences, including thermophilic homologues, were completely excluded from THOR pretraining. We selected five bacterial MDH lineages spanning divergent regions of the MDH phylogeny (Fig. 3A; Methods M7), with an average aligned pairwise sequence identity of only 36% between the starting enzymes (Supplementary Fig. 16). Natural thermophilic MDHs occupied phylogenetically distinct lineages from the selected mesophilic starting sequences, rather than forming obvious matched mesophile–thermophile pairs around each wild-type background (Supplementary Fig. 17). Together with the substantial sequence divergence with an average maximum pairwise sequence identity of 52% (Supplementary Fig. 18), this means that no obvious one-to-one evolutionary correspondence exists between the selected mesophilic starting sequences and natural thermophilic MDHs, providing a stringent test of whether thermal-adaptation information can be transferred without matched homologous examples.

**Figure 3.**
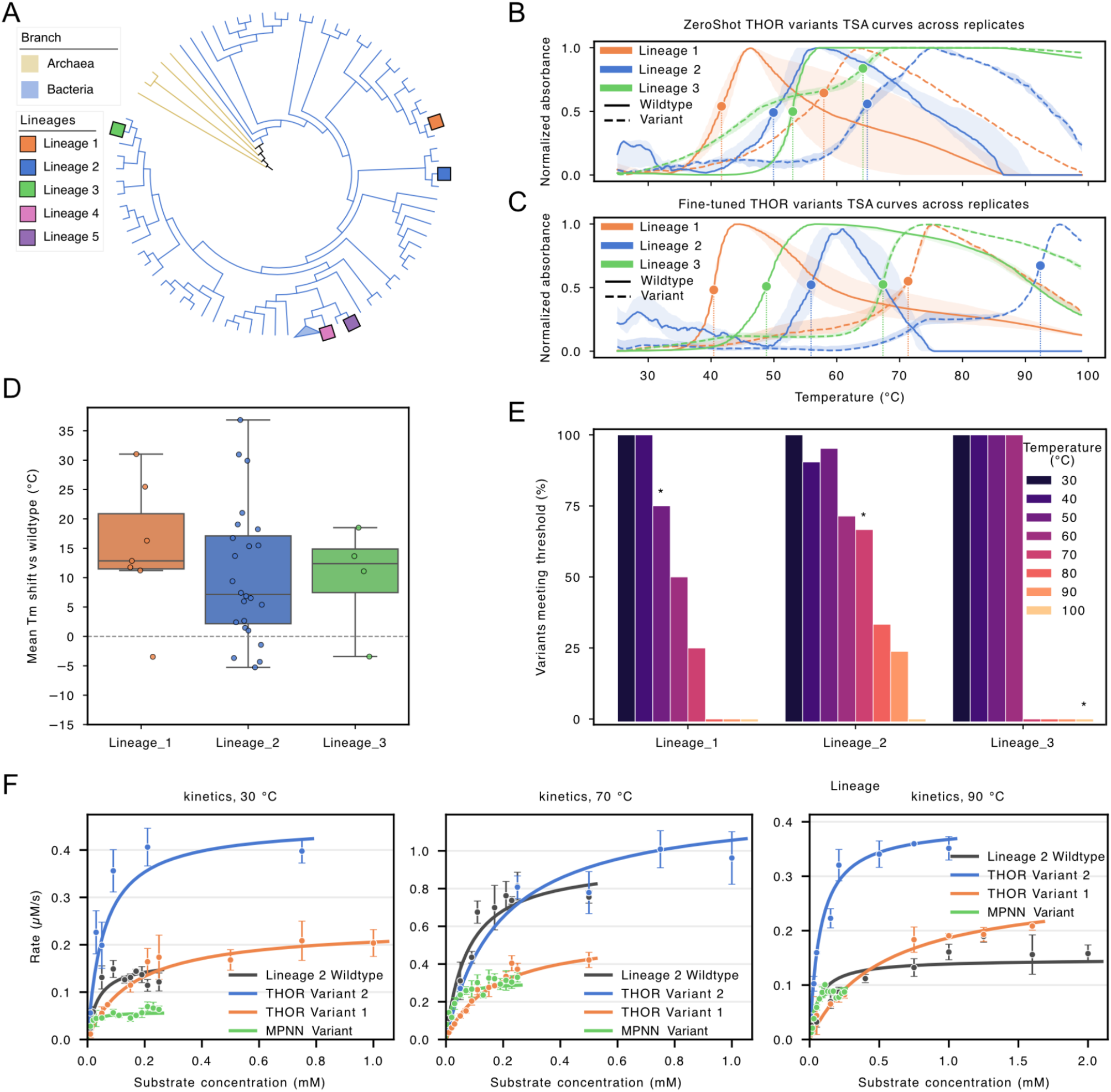
| THOR increases thermal robustness across divergent malate dehydrogenase lineages while retaining catalytic activity. **A**, Representative phylogenetic tree of malate dehydrogenase sequence diversity, showing the lineages selected for experimental validation. **B,** Thermal denaturation profiles of the top-performing zero-shot generated THOR variant from each of three divergent MDH lineages and the corresponding wild types. Shaded areas represent replicate range. **C,** Thermal denaturation profiles of the top-performing enzymes after fine- tuning. **D,** Change in apparent melting temperature (*T_m_*) of THOR variants relative to their corresponding wild types across the three lineages. **E**, Fraction of enzymes retaining detectable activity following thermal challenge at increasing temperatures, showing lineage-dependent extension of the active temperature range. * indicates WT enzyme. **F**, Steady-state enzyme kinetics for Lineage 1 wild type, two THOR variants and a ProteinMPNN-designed variant measured at 30, 70 and 90 °C. THOR variants combine increased thermal robustness with retained, and in some conditions enhanced, catalytic activity, whereas increased stability alone does not necessarily preserve catalytic function (details of kinetics parameters in Supplementary Table 3).

We first used the pretrained model to generate thermally adapted variants from Lineage 1-3 MDH starting sequence (3 per family), providing a zero-shot test of cross-family transfer. Each of the three lineages yielded active variants suitable for subsequent characterisation (Supplementary Fig. 19). Apparent melting temperatures were measured by differential scanning fluorimetry (Methods M8, Supplementary Data File 1). Although MDH was entirely absent from pretraining, THOR-generated variants showed significantly increased (median shift of +10.9 °C) apparent *T_m_* relative to their corresponding wild types (stratified Wilcoxon rank-sum test, p-value = 0.029), with the best design melting at 65.5 ± 1.3 °C versus 50.1 ± 1.0 °C for its wild type (mean ± SD, n = 3), an increase of 15.4 °C, (Fig. 3 B, Supplementary Fig. 20), demonstrating that thermal-adaptation information learned outside the MDH family could produce an experimentally measurable thermal shift in a new evolutionary context.

We next asked whether limited MDH-specific information could further increase this shift. Starting from the pretrained model, we fine-tuned THOR on mesophilic and thermophilic MDH homologues, without imposing one-to-one sequence correspondences between the two groups. Only 67 thermophilic MDHs were available (Supplementary Fig. 21), and the closest examples shared only 72.4% sequence identity with the selected starting sequences and occupied distinct thermophilic lineages (Supplementary Fig. 17,18). Following fine-tuning (Methods M9), we generated an expanded set of 39 designs across all five wild-type lineages. All 27 designs from Lineages 1–3 retained enzymatic activity, whereas none of the four designs from Lineage 4 exhibited detectable activity, including the wild type. Of the eight designs from Lineage 5, only three could be successfully expressed and purified, and these exhibited only marginal activity relative to their wild type (Supplementary Fig. 22). Consequently, subsequent analyses focused on the 27 active designs from Lineages 1–3. 85% of designs (n=23) assayed by thermal shift, THOR-generated variants showed significantly increased apparent *T_m_* relative to their corresponding wild types (stratified Wilcoxon rank-sum, p-value = 0.0013; median shift +13.7 °C, 95 % CI 8.2–18.8 °C), the strongest design in each lineage increasing *T_m_*by 31.0 °C, 36.8 °C and 18.5 °C respectively (Fig. 3C); the best overall, Lineage 2 Variant 1, melted at 92.3 ± 0.2 °C versus 55.5 ± 1.6 °C for its wild type (mean ± SD, n = 3 wells). More broadly, 70 % (19 of 27 active tested variants) increased *T_m_* by at least 5 °C relative to their corresponding wild type, with positive thermal shifts observed across all three lineages (Fig. 3D, Supplementary Fig. 23, 24) Together, these results indicate that THOR variants preserve a functional folded state while shifting the thermal limits of the enzyme.

We next tested whether the increases in apparent *T_m_* were accompanied by improved retention of activity after thermal stress. THOR variants were incubated for 10 min at increasing temperatures, cooled, and assayed for MDH activity (Methods M8). In lineages 1 to 3, 74% of variants retained measurable activity after exposure to temperatures above the apparent *T_m_* of the corresponding wild type, with several remaining active after treatment at 80-90 °C (Fig. 3E, Supplementary Fig. 25). The wild type of Lineage 3 likewise showed activity far beyond its apparent *T_m_*, the wild-type enzyme recovered activity after heating across all of the tested temperature range. Thus, apparent *T_m_* and post-heating activity captured related but distinct aspects of thermal behaviour, consistent with their moderate correlation across variants (Spearman’s *ρ* = 0.69, p-value = 6.3e-5; Supplementary Fig. 26). Across divergent MDH backgrounds, THOR therefore improved both the apparent unfolding transition and the ability of the enzyme to remain functional after thermal stress.

To test whether the large thermal gains in MDH reflected an unusually permissive scaffold rather than a THOR-specific effect, we benchmarked THOR against ProteinMPNN, a widely used inverse-folding model^20^. Unlike THOR, ProteinMPNN does not explicitly learn sequence features associated with thermal adaptation, but instead identifies sequences compatible with a specified protein backbone. We selected the lineage 2 wild type, for which THOR produced the largest increases in apparent *T_m_* and strong catalytic activity, and redesigned the same sequence using the ProteinMPNN protocol as reported for generating stable enzyme variants preserving the active site^21^. Following the protocol, four ProteinMPNN sequences resulting in sequences spanning 53%–83% sequence identity to the wild type were generated and expressed in *E. coli* using the same workflow as the THOR variants. All four ProteinMPNN variants expressed successfully, with apparent melting temperatures ranged from 46 to 74 °C, corresponding to increases of up to 18 °C relative to the wild type (Supplementary Fig. 27). However, only one of the four variants retained detectable MDH activity (Supplementary Fig. 28). This is consistent with recent reports showing that inverse-folding designs can improve foldability or thermal stability without necessarily preserving catalytic function^31–33^. Thus, generic structure-conditioned redesign was sufficient to increase apparent *T_m_*in MDH, but this did not consistently preserve catalytic function. In contrast, THOR transferred the thermal phenotype while retaining activity across multiple variants, supporting the idea that the model captures adaptation-associated sequence information rather than simply favouring sequences compatible with a stable fold.

### Thermostability can be increased without compromising catalytic turnover

To determine whether the large thermal shifts produced by THOR remained compatible with catalytic performance, we selected the two lineage 2 variants with the largest increases in *T_m_*, variants 2 and 1 (+31 °C and +36 °C relative to wild type; Fig 3C), and measured Michaelis–Menten kinetics at 30 °C, 70 °C and 90 °C (Supplementary Table 3, Methods M8) with wild-type MDH and the active ProteinMPNN-designed variant included as controls. Variant 2 exceeded wild-type turnover at all three temperatures, with *k_cat_* values of 2.71×, 1.32× and 2.67× wild type at 30 °C, 70 °C and 90 °C, respectively. Variant 1 also exceeded wild type at 30 °C and 90 °C (1.42× and 1.91×, respectively), although its turnover was reduced at 70 °C. By contrast, the single active ProteinMPNN control showed an 83–92% reduction in *k_cat_* relative to wild type across the same temperature range. The large increases in apparent *T_m_* achieved by THOR therefore remained compatible with high catalytic turnover from mesophilic to high-temperature conditions. Although *K_M_* values were not improved to the extent of *k_cat_* (Supplementary Table 3), selected THOR variants reached almost threefold higher *k_cat_* than wild type, showing that substantial thermal gains can coexist with high, and markedly enhanced, catalytic turnover.

### Distinct patterns of protein dynamics converge on enhanced thermostability

Thermostability can arise through different combinations of structural and dynamic changes rather than a single universal mechanism, and excessive rigidification can be detrimental to enzyme function ^34–36^. Because THOR introduced different sequence changes across divergent MDH backgrounds (Fig. 2C, D, Supplementary Fig. 10,11) while producing similar thermal phenotypes (Fig. 2B, 3D), we asked whether these variants converged on a common change in protein dynamics or reached increased thermal stability through distinct, background-dependent dynamical responses.

We first compared the structures of the selected wild-type MDHs. Despite sharing only 36% average pairwise sequence identity, the enzymes adopted highly similar folds, with an average TM-score above 0.95 (Supplementary Fig. 29). We therefore examined whether the thermal gains introduced by THOR were associated with changes in local protein dynamics. We performed 300-ns molecular dynamics simulations (4 replicates per variant, Methods M10) for all active THOR variants with experimentally validated *T_m_* increases of at least 5 °C relative to their corresponding wild type and quantified residue-level changes in flexibility as change in Root Mean Squared Error (ΔRMSF) (Fig. 4A). Across all three lineages, THOR variants showed spatially localised regions of both increased and decreased mobility, consistent with previous MD studies demonstrating that thermostabilization can involve heterogeneous changes in protein dynamics rather than uniform rigidification^3735^.

**Figure 4.**
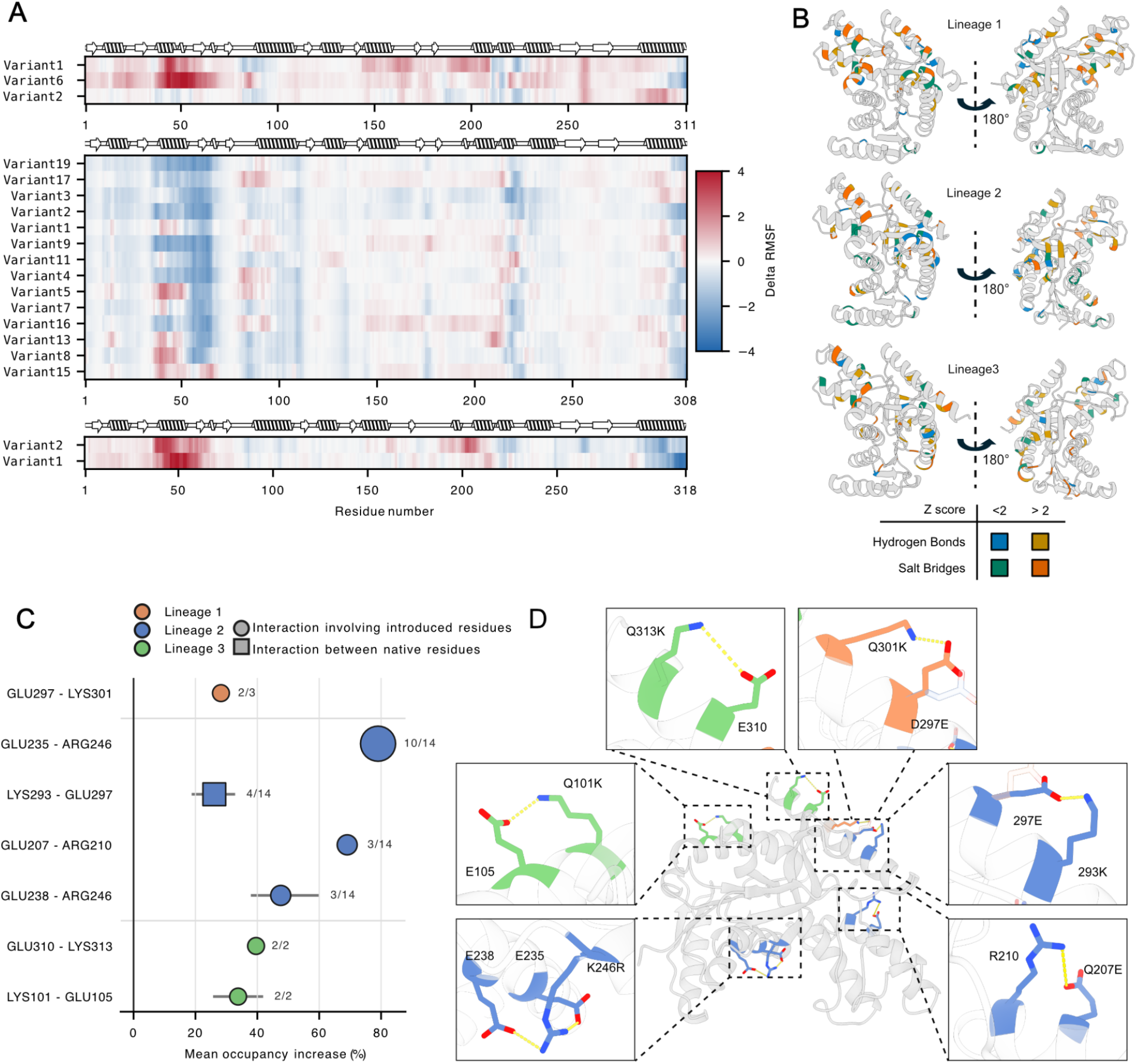
| THOR variants realise thermal adaptation through distinct changes in protein dynamics and intramolecular interactions. A, Residue-level changes in root-mean-square fluctuations (ΔRMSF) of THOR variants relative to their corresponding wild types across three MDH lineages (average of 4 simulation results is per variant, Methods M10). Secondary-structure elements are shown above each heatmap (Methods M10). **B,** Structural mapping of hydrogen-bond and salt-bridge changes in representative variants from each lineage. Residues showing the strongest changes in interaction occupancy are highlighted on two views of each structure. **C,** Interactions showing increased occupancy across THOR variants, grouped by lineage. Symbol size indicates the magnitude of the occupancy increase, while symbol shape distinguishes interactions involving an introduced mutation (circle) from those arising through conformational rearrangement of native residues (square); labels indicate the number of variants in which each interaction was observed. **D,** Representative structural examples of newly introduced or strengthened hydrogen-bond and salt-bridge interactions associated with THOR substitutions.

Moreover variants derived from the same wild-type background differed in which regions became more or less mobile, i.e. within lineage 2, variants derived from the same wild-type background showed distinct dynamical responses: pairwise ΔRMSF profiles were only moderately correlated (median Pearson’s *r* = 0.51), range -0.43 to 0.88), and one third of strongly affected residue positions (|Z| > 2 in at least one variant; 25 of 75) showed opposite changes in mobility across different variants. The direction of these changes differed between backgrounds: lineages 1 and 3 were biased towards increased mobility (4.4% and 6.1% of residue observations with Z > 2, versus 2.1% and 2.4% with Z < −2), whereas lineage 2 was dominated by reduced mobility (5.9% with Z < −2 versus 1.8% with Z > 2) (Supplementary Fig. 30)

Mapping these changes onto the structures showed that the affected regions were distributed differently across the three lineages (Fig. 4B). Variants gained salt bridges relative to their wild type in all three backgrounds (17 of 19 variants above WT; pooled mean +1.33, Wilcoxon rank-sum test, p-value = 2.7 × 10⁻⁵), with no difference in the size of this gain between lineages (Kruskal-Wallis test, p-value = 0.40). Hydrogen bonding showed no overall shift (pooled mean +0.28, p-value = 0.44) but differed between backgrounds (p-value = 0.024): lineage 1 variants lost hydrogen bonds (−4.5, p-value = 0.014) and residue contacts (−16.3, p-value = 0.008), whereas lineages 2 and 3 gained both. Increased apparent *T_m_* was therefore associated with a consistent enrichment of salt bridges superimposed on background-specific redistributions of local dynamics and hydrogen bonding (Supplementary Fig. 31).

We next examined how these distinct dynamical responses were reflected in local interaction networks. THOR variants strengthened different combinations of hydrogen bonds and salt bridges in each background (Methods M10), i.e. lineage 2 showed the broadest set of recurrent changes, including E235–R246 in 10 of 14 variants, K293–E297 in 4 of 14, and E207–R210 and E238–R246 in 3 of 14 variants each. By contrast, lineage 1 was dominated by E297–K301 (2/3 variants), whereas lineage 3 consistently strengthened E310–K313 and K101–E105 (2/2 variants each) (Fig. 4C). Structural examples show that most strengthened interactions directly involve introduced residues, whereas others, such as K293–E297, strengthen between residues already present in the wild type (Fig. 4D). Together, these simulations show that similar thermal gains can arise through distinct local dynamical and interaction changes, indicating that transferred thermal adaptation is implemented in a background-dependent manner rather than through a single conserved stabilisation mechanism.

## Discussion

Thermostability has evolved independently in diverse protein families, often without changing their enzymatic function. The key question is whether the sequence features responsible for thermal adaptation can be separated from the evolutionary history of each protein family. This distinction matters both conceptually and practically: if adaptive information can be separated from its original evolutionary context, it may be transferable to proteins that never experienced the same evolutionary trajectory, reducing the need to rediscover the same property independently within each protein family. Our results provide experimental evidence that thermal adaptation contains such a transferable component. Information learned across diverse mesophilic and thermophilic enzymes produced substantial thermal shifts in MDH despite the complete absence of MDH from training, with zero-shot variants increasing apparent *T*_m_ by up to ∼15 °C (Fig. 3B). Limited subsequent supervision to thermophilic MDHs extended these shifts to up to ∼36 °C (Fig. 3C). Thermal adaptation is therefore not confined to family-specific evolutionary solutions, but contains information that can be learned across distant proteins and redeployed in a new sequence context.

The value of such transfer is amplified by the way thermostability information is distributed across protein sequence space. Large-scale thermal proteome datasets provide experimentally measured melting temperatures, while mutation databases catalogue stability effects of individual substitutions, but together they cover only a small fraction of known protein diversity^9,19,38–40^. Organism growth temperature provides much broader coverage^6^, but is an indirect proxy for protein thermostability and is strongly biased towards mesophilic organisms. Crucially, thermophilic examples are sparse within individual protein families (Supplementary Fig. 2), making it difficult to separate temperature-associated sequence features from phylogeny and family-specific constraints using conventional supervised learning. THOR exploits a different source of information: differences between unpaired mesophilic and thermophilic sequence populations distributed across many unrelated families. In this setting, the adaptive state of the population itself provides supervision, without requiring measured thermostability examples or known evolutionary transformations within the target family.

THOR differs from both experimental and computational routes to thermostability engineering. Directed evolution, deep mutational scanning and combinatorial mutagenesis explore variants around an existing protein and therefore remain anchored to the evolutionary solution represented by that starting sequence^13,14,41^. Because functional protein sequence space is sparse and highly structured^42,43^, such searches sample only a small subset of the stabilising solutions explored across evolution. Ancestral sequence reconstruction exploits evolutionary history to access sequence states beyond extant proteins, but by construction remains tied to the phylogeny of the protein family being reconstructed ^44,45^. Physics-based and structure-guided approaches can identify mechanistic routes to stabilisation, while inverse-folding and multimodal models generate sequences compatible with structural and functional constraints^20,46,47^. Supervised models can further predict or rank *T*_m_, Δ*T*_m_ or ΔΔG from labelled examples^48–52^. These approaches are powerful, but none is explicitly designed to isolate a phenotype-specific component of adaptation from the broader evolutionary context in which it arose. The distinction here is therefore not simply access to a larger sequence space, but the ability to learn an adaptive signal across evolutionary lineages and test whether it remains meaningful outside those lineages. The transferable component of thermal adaptation does not appear to correspond to a fixed thermophilic sequence programme. Natural thermophiles show recurring compositional and substitutional biases across evolutionary lineages^7,8,53^, and THOR recovered part of this signal across held-out protein families. Yet generated sequences did not converge on a common thermophilic solution. Different starting proteins acquired distinct substitutions while retaining their underlying sequence and phylogenetic context. Thermal adaptation therefore appears many-to-one at the sequence level: similar thermal phenotypes can be reached through different mutational routes (Fig. 2C–F). What transfers across evolution may therefore be shared sequence features of the thermophilic state, rather than a single sequence solution. More broadly, distinct evolutionary states may provide enough information to learn transferable directions in sequence space even when direct molecular measurements are sparse.

Importantly, the transferred phenotype extended beyond an increase in apparent *T_m_*. THOR variants retained activity after heat challenge, and selected variants maintained high catalytic turnover across temperatures from 30 °C to 90 °C (Fig. 3C–E). To test whether MDH simply tolerates extensive stabilising redesign, we generated four ProteinMPNN variants from the same wild-type lineage. All expressed successfully and several showed increased apparent *T*_m_, yet only one retained detectable MDH activity (Supplementary Fig. 28), indicating that increased stability in this scaffold is not generally achieved without functional cost. Selected THOR variants reached nearly threefold higher *k_cat_* than the wild type while also showing among the largest increases in apparent *T_m_* (Fig. 3E). Although *K_M_* and *k_cat_*/*K_M_* were not consistently improved (Supplementary Table 3), the data show that substantial gains in thermal robustness can be compatible with and accompanied by increased catalytic turnover.

At the molecular level, similar thermal phenotypes were realised through different physical routes. Although the three MDH backgrounds retain highly similar global folds (Supplementary Figure 29), THOR variants showed markedly different redistributions of local dynamics, with some lineages dominated by reduced mobility and others by increased mobility (Fig. 4A). Even variants derived from the same wild-type background altered different structural regions, with similarly distinct changes in hydrogen bonding and salt-bridge interactions (Fig. 4A, C). This argues against global rigidification as a universal mechanism of thermal adaptation, despite its frequent association with thermostability^54–56^, and is consistent with evidence that high thermal stability can arise through distinct structural and dynamical solutions^35,36,57^. The sequence and molecular dynamics analyses therefore suggest that thermal adaptation can be transferred without preserving a common sequence or dynamical mechanism.

More broadly, repeated evolution may contain information about adaptation that extends beyond the individual solutions realised within particular lineages. Experimental validation here is centred on MDH, the largest thermal shifts follow limited family-specific refinement, and broader testing will be required to establish how generally this principle applies across folds and catalytic mechanisms. Nevertheless, the zero-shot results from non-homologous thermostability transfer show that thermal adaptation signal generalises across distant evolutionary backgrounds. Whether analogous adaptive axes can be extracted for properties such as pH, solvent or other environmental tolerance remains an important question. Evolution may therefore encode transferable principles of adaptation beyond the specific sequence solutions found in individual lineages, allowing them to be transferred into molecular contexts in which they never evolved.

## Acknowledgments

This study was supported by the Biotechnology and Biological Sciences Research Council (BBSRC; grant BB/Y000730/1), the Swedish Research Council (Vetenskapsrådet; grants 2023-04254 and 2019-05356), Formas (grant 2022-02316), the Knut and Alice Wallenberg Foundation through the WASP-DDLS postdoctoral programme, and the Marius Jakulis Jason Foundation scholarship. Computational resources and data handling were provided by the National Academic Infrastructure for Supercomputing in Sweden (NAISS), partly funded by the Swedish Research Council through grant agreement 2022-06725, and by the Berzelius resource at the National Supercomputer Centre (NSC), funded by the Knut and Alice Wallenberg Foundation. The authors also acknowledge the use of resources provided by the Isambard-AI National AI Research Resource (AIRR). Isambard-AI is operated by the University of Bristol and funded by the UK Government’s Department for Science, Innovation and Technology (DSIT) through UK Research and Innovation and the Science and Technology Facilities Council [ST/AIRR/I-A-I/1023].

## Methods

### M1 Data collection and preprocessing

#### Environmental-temperature data for regression

The environmental temperature (ET) dataset^58^ was used to train the pET regression model. To reduce sequence redundancy, sequences were first clustered at 95% sequence identity using CD-HIT^59^ . Sequences longer than 512 amino acids were excluded to standardize the maximum input length. The resulting non-redundant ET dataset was subsequently clustered at 30% sequence identity using MMseqs2^60^. An independent test set was generated by randomly sampling approximately 2.3% of the filtered dataset while preserving the temperature distribution of the full dataset. This resulted in a test set containing 62,930 sequences, with the remaining sequences retained for model training. To evaluate performance across the temperature range without over-representing densely sampled intervals, the test set was further subsampled to obtain an approximately uniform temperature distribution, yielding a balanced subset of 636 sequences.

#### Sequence populations for THOR pretraining

THOR was pretrained on separate mesophilic and thermophilic sequence populations and subsequently fine-tuned on MDH sequences as described in M9. The mesophilic population was derived from 16,044,844 UniProt enzyme sequences. The thermophilic population was derived from 1,098 metagenomic samples collected from environments above 60 °C and downloaded from JGI/IMG^61^, comprising 51,337,401 sequences before filtering. Both populations were filtered to retain sequences of 50–512 amino acids and clustered at 100% sequence identity using CD-HIT to remove duplicates. Sequences were annotated with Clusters of Orthologous Groups (COG) identifiers and Enzyme Commission (EC) numbers using eggNOG-mapper^62^; sequences lacking either annotation were excluded. After filtering and annotation, the dataset contained 10,651,518 mesophilic and 6,603,626 thermophilic sequences. Ten root-level COG annotations were selected for family-level evaluation, and all sequences assigned to these COGs were removed from the pretraining dataset. This held-out set contained 340,037 mesophilic and 110,215 thermophilic sequences. These sequences were not used for parameter optimisation, but were used to monitor convergence and evaluate the model. The selected COGs and their sequence counts are listed in Supplementary Table 2.

### M2 Regression model Training and evaluation

ET labels were centred by subtracting the midpoint between the minimum and maximum ET values in the dataset. For training, the ET range was divided into ten equal-width temperature bins, and sequences were sampled uniformly across bins to balance temperature representation.

The regression model used a modified deep convolutional architecture^51^ with a self-attention layer and a final dense output layer. Three models with identical architectures and training and validation splits were trained from independent parameter initialisations. Each model minimised the mean squared error (MSE) between predicted and reported ET values. Models were trained for up to 100 epochs, with a batch size of 256, using the Adam^63^ optimiser at a learning rate of 0.001. The checkpoint with the lowest validation MSE was retained for each model.

For inference, pET was calculated as the mean prediction of the three models. Predictions were converted back to the original temperature scale before evaluation. Performance was assessed on both the full test set and its approximately temperature-balanced subset using the coefficient of determination (R²) and mean absolute error (MAE).

### M3 THOR Model Architecture and Training

The THOR models were designed to utilize additional probability features alongside sequence data for training. These probability features were computed using the ESM1v^64^ model by sequentially masking 10% of the input sequence at a time and predicting the amino acid probabilities at each masked position. This process was repeated iteratively until probabilities for all sequence positions were obtained. Both mesophilic and thermophilic sequences in the finetuning and pretraining datasets had these features precomputed prior to training. Additionally, all sequences were padded to a maximum length of 512 amino acids to ensure consistent input dimensions for the model.

The generative model in THOR is trained using a pair of generators G, F, and corresponding discriminators Dx,Dy. The generators are based on modified U-Net architectures with attention gating^65,66^ using ∼75 million parameters. The discriminator architecture is patch-based convolutional models^67^ that include two self-attention^68^ layers and spectral normalization^69^ with 33 million parameters. Building on the original adversarial and cycle-consistency objectives^27^, we introduce a novel evolution-guided loss *L_evo_* that incorporates residue-specific evolutionary constraints directly into sequence generation. Using ESM1v-derived amino-acid likelihoods to weight position-wise cross-entropy between input (*x_i_*) and generated (*x_i_’*) sequences, this objective preferentially preserves evolutionarily supported residues^28^ while enabling sequence exploration for thermal adaptation.

Here, CE denotes cross-entropy, *x_i_* denotes the input residue at position *i*, and *x_i_’* denotes the corresponding generator output, written as *G(x_i_)* in the equation. The scalar *l_i_* is the ESM1v-derived probability of the input residue at that position. The corresponding term for *F* applies to the reverse transformation.

Thus, the total loss function *L(G, F, D_X_, D_Y_)* for training combines multiple components:

1. Adversarial Loss (*L_GAN_*): Encourages the generators to produce sequences indistinguishable from the real data distributions of mesophiles and thermophiles.
2. Cycle Consistency Loss *(L_cyc_)*: Ensures the cyclic mapping between mesophilic and thermophilic sequences preserves the original sequence characteristics.
3. Evolutionary Loss *(L_evo_)*: Enforces evolutionary constraints as described above.

The combined loss function is expressed as:

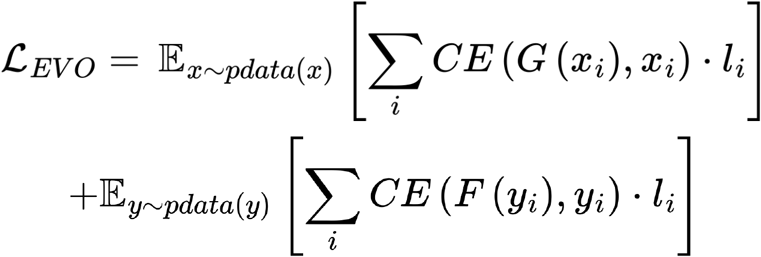

where λ is a scaling parameter that balances the influence of *L_evo_* relative to other loss components, during training, λ is dynamically adjusted using a PID controller to maintain the desired median sequence identity between the generated and input sequences.

The adversarial loss is defined as: L_GAN_(G,DY,X,Y)=E_y∼pdata(y)_[logD_Y_(y)]+E_x∼pdata(x)_[log(1−D_Y_(G(x)))]

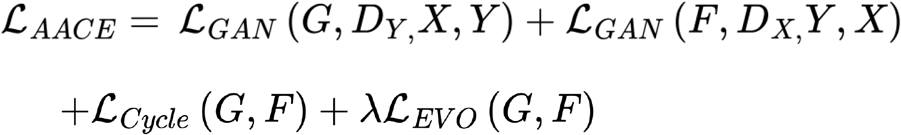

and the cycle consistency loss is defined as:

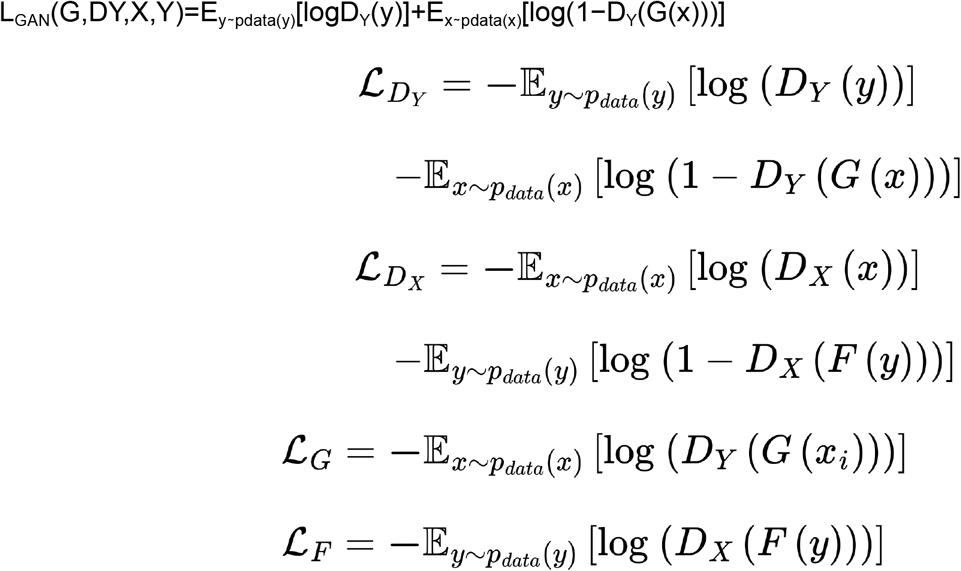

A schematic representation of the training setup, including data flow and loss components, is provided in Supplementary Figs. 4–6. Briefly, THOR was trained using an unpaired adversarial cycle-consistent framework^27^, incorporating additional evolutionary consistency and sequence-divergence constraints to preserve the identity of the wild-type protein. Training was performed on unpaired mesophilic and thermophilic enzyme sequences represented using ESM-derived features, with a maximum sequence length of 512 residues. The generators and discriminators comprised convolutional architectures with five and three filter stages, respectively. The training objective combined adversarial, cycle-consistency, evolutionary consistency and sequence-preservation losses, with corresponding weights of 1.0 for cycle consistency, 0.5 for each evolutionary consistency term and 0.85 for self-consistency. Sequence divergence was dynamically controlled using a proportional–integral–derivative (PID) controller with coefficients K_p_=5, K_i_=0.5 and K_d_=0.1. The generators and discriminators were optimised separately using Adam^63^, with learning rates of 2x10^-5^ and 1x10^-5^, respectively, and β_1_=0.5 and β_2_=0.99. Models were trained with a batch size of 512 for up to 30,000 steps, using 10,000 warm-up steps followed by cosine learning-rate decay. Validation was performed every 100 training steps. Training typically converged within 48 hours using four NVIDIA A100 GPUs.

### M4 Amino acid Association and Substitution analysis

#### Amino acid composition and enrichment

Analyses were carried out on the ten held-out test families, comprising 340,037 mesophilic sequences from UniProt and 110,215 thermophilic sequences from IMG. Amino acid frequencies were computed separately for each group by counting occurrences of each of the 20 standard residues and normalising by the total number of standard residues in that group; non-standard residues were excluded. Variants were generated with a pretrained THOR model for every mesophilic test sequence of 100–512 residues with a predicted ET of at most 45 °C (260,609 sequences). Two variants were sampled per wild type and the one with the higher predicted ET was retained; retained variants shared on average 84% sequence identity with their wild type (range 75–95%). Variant amino acid frequencies were computed as for the wild-type groups. Enrichment relative to the mesophilic group was expressed as log₂ fold change and tested for each amino acid with a two-sided Fisher’s exact test on a 2 × 2 table of that residue against all other standard residues, with Benjamini–Hochberg correction across all amino acids and both comparisons; residues with an adjusted P below 0.01 are marked.

#### Temperature-associated substitutions in natural sequences

Each of the ten test families was clustered separately with MMseqs2 at 30% sequence identity (easy-cluster --min-seq-id 0.3). Clusters with at least ten members containing both mesophilic and thermophilic sequences were retained and aligned with Clustal Omega^70^, giving 60 alignments. Alignments with fewer than five sequences in either group were excluded, leaving 43. Following Pinney et al.^11^, every combination of alignment position and amino acid was tested for temperature association with a two-sided Fisher’s exact test comparing presence of that residue between the two groups, with Benjamini–Hochberg correction across all tests within an alignment at α = 0.01. Positions carrying at least one mesophile-associated and one thermophile-associated residue were retained; 27 alignments, drawn from seven of the ten families, contained such positions. Each retained position contributed to a 20 × 20 substitution matrix weighted by the product of the group-specific frequencies of its associated residues, normalised so that every informative position contributed equally. Contributions were summed across alignments and converted to relative frequencies. For each of the 190 unordered residue pairs, the preferred direction was assessed with a two-sided binomial test (α = 0.05) on the per-position directional contributions, restricted to pairs supported by at least 100 informative positions and corrected with Benjamini–Hochberg at *α* = 0.01; 41 pairs showed a significant preferred direction.

#### Substitutions introduced by THOR

Each residue of a variant was compared with the residue at the same position of its wild type, giving 14,951,483 substitutions across the 260,609 wild-type–variant pairs. Unchanged positions and non-standard residues were excluded, and counts were normalised to relative frequencies. The overlay was matched to the natural matrix: the same number of pairs (41) was marked, taken in order of the frequency of the preferred direction, one direction per unordered pair. Marks in the two panels consequently have different meanings, statistical significance for the natural matrix, rank by frequency for the model, and the boundary of the ranked set falls between pairs of similar frequency (89,408 and 89,093 substitutions). Of the 41 pairs marked in each panel, 24 are marked in both, and in all 24 the model’s preferred direction agrees with the natural one. Comparison of the two matrices. The pooled natural and model matrices were compared across all 380 directed amino acid changes (20 × 19), excluding the unchanged diagonal and retaining changes with zero frequency in either matrix; no filter on frequency rank or preferred direction was applied. Agreement was quantified with Pearson (*r* = 0.80, nominal p-value = 1.8 × 10⁻⁸⁵) and Spearman (ρ = 0.75, nominal p-value = 1.3 × 10⁻⁷⁰) correlation. Substitution frequencies are compositional and the 380 categories are not independent, so these nominal P values do not account for dependence between substitution types, sequences or families.

### M5 U-map analysis of embeddings

Embeddings for all wild-type sequences from the 10 test families, as well as their corresponding variants generated by the pretrained THOR model, were computed using ESM1b^71^. To visualize the relationships between these sequences, the embeddings of both wild-type and variant sequences were projected to two dimensions using UMAP^30^ with cosine similarity as distance metric.

### M6 Phylogenetic Tree analysis

For each of ten test set COG families, we aligned natural mesophiles, natural thermophiles, one designed variant per wild type and outgroup sequences with MAFFT-DASH^72^. We trimmed the alignment with ClipKIT^73^ and inferred a maximum-likelihood tree with IQ-TREE^74^ (30,235 tips in total). A natural sequence was classed as thermophilic if its source environmental temperature was ≥ 60 °C. THOR-predicted temperatures were thresholded 60 °C cutoff was used only as a robustness check. 94 of 773 natural thermophiles are longer than regression models 512-residue limit. They were not truncated: they stayed in the trees but were left out of every analysis that uses predicted values. Each tree was rooted on the outgroup, or at the midpoint where no outgroup was retained. It was then split into disjoint monophyletic clades of at most 75 natural tips, and each designed variant was assigned to its wild type’s clade. For each clade we computed two outcomes: the fraction of variants predicted at or above threshold; the median gain in predicted temperature of each variant over its wild type (ΔT).

Clades containing a natural thermophile (n = 147) were compared with clades containing none (n = 297), with clades as the unit of replication. Differences were tested by permuting design–wild-type assignments on the fixed tree (10,000 permutations), with confidence intervals from bootstrapping clades. They were also adjusted, by weighted least squares, for the mean predicted temperature of each clade’s wild types and for log clade size. Because the hypothesis of interest is one of no difference, we tested equivalence by two one-sided tests. The margins, fixed in advance, were ±10 percentage points for the hot-variant fraction and ±5 °C for ΔT. Families were analysed separately and combined by Stouffer’s method, weighted by the square root of clade count. COG4188 has no natural thermophiles and is reported separately.

To test whether designs move toward natural thermophiles in sequence space, we used IQ-TREE pairwise ML distances. For each wild-type/variant pair we computed the change in rank of the nearest natural thermophile among natural sequences, excluding the pair’s own wild type. The secondary statistic was the change in thermophile fraction among the ten nearest natural neighbours. We used ranks because they are unaffected both by the uniform increase in distance from about 30 substitutions per design and by ML saturation. Specificity was checked against ten size-matched random sets of natural mesophiles per family. Significance was tested by a Wilcoxon signed-rank test on each clade’s median Δrank. Full details, including the alternative partition criteria and the reasons for these choices, are in Supplementary Methods.

### M7 Generating and Filtering Variants for Experimental Validation

For zero-shot design with pretrained THOR, two MDH sequences were selected from UniProt^26^ and one from MGnify^75^ as wild-type queries. Query sequences shared at most 80% sequence identity with the training sequences. For each query, 2,000 variants were generated. Each proposed substitution was evaluated using the ESM1v probability features described in M3, and substitutions with probabilities below 0.05 were reverted to the wild-type residue. The filtered variants were ranked by pET using the regression model (M2), and the three highest-ranked variants per wild type were selected for experimental validation. The selected sequences are provided in Supplementary Data File 1. Fine-tuning and the expanded MDH design set are described in M9.

### M8 Enzymatic activity assay and kinetics, melting temperature measurements

#### Apparent melting temperature

Expressed and purified wild-type and engineered MDHs were stored at 20–50 µM in 300 mM NaCl and 50 mM NaH₂PO₄ (pH 7.4). Apparent melting temperatures (Tm) were measured by differential scanning fluorimetry using SYPRO Orange. Reactions contained 3–5 µM protein and 5× SYPRO Orange in the same buffer, in a final volume of 20 µl, and were assembled in white 96-well qPCR plates. Buffer replaced protein in blank controls. Measurements were performed in triplicate on an Mx3005P qPCR instrument with a temperature ramp of 1 °C/min and excitation and emission filters of 492 and 620 nm, respectively.

#### MDH activity assay

Activity was measured by monitoring NADH oxidation as a decrease in absorbance at 340 nm (A340) using a FLUOstar Omega microplate reader (BMG Labtech). Enzyme stocks (1 µg/µl) were diluted to 25 µg/ml in 20 mM HEPES (pH 7.4). NADH (1.5 mM) and oxaloacetate (OAA; 2 mM) solutions were freshly prepared in the same buffer. Mixtures containing 50 µl enzyme, 50 µl NADH and 150 µl HEPES buffer were pre-incubated in a thermomixer for 10 min at 30–80 °C in 10 °C increments. Reactions were initiated by adding 50 µl OAA, giving a total volume of 300 µl. A340 was recorded every 30 s for 10 min. Conditions were assayed in triplicate, with enzyme-free blanks containing NADH, OAA and buffer. Activity was calculated from the initial linear decrease in A340 after blank subtraction and expressed relative to the 30 °C control.

#### Michaelis–Menten kinetics

Initial rates were measured for wild-type MDH, Variant 1, Variant 2 and MPNN70 over an OAA concentration range of 0.01–1.2 mM, adjusted for each variant, at a fixed, near-saturating NADH concentration. Measurements used the FLUOstar Omega reader at 30 °C in 20 mM HEPES (pH 7.4). Enzyme concentrations were 0.5 ng/µl for wild type and THOR variants and 2 ng/µl for MPNN70. NADH consumption rates (mM/s) were calculated from the decrease in A340 using an extinction coefficient of 6.22 mM⁻¹ cm⁻¹. Each condition was measured in triplicate. Initial velocities were fitted by nonlinear regression to the Michaelis–Menten equation, *V = k_cat_[E][S]/(K_m_+[S])*, to estimate *K_m_* and *k_cat_*; catalytic efficiency was calculated as *k_cat_/K_m_*. Here, [E] and [S] denote enzyme and substrate concentrations, respectively.

### M9 Fine tuning THOR

The fine-tuning dataset was assembled from the non-redundant ET dataset. Sequences were annotated with COG and EC identifiers using eggNOG-mapper, and sequences assigned to both COG0039 and EC 1.1.1.37 were retained. Partitioning by ET yielded 3,486 mesophilic and 67 thermophilic sequences.

Fine-tuning resumed from the pretrained THOR checkpoint, including the saved optimiser and PID-controller states, using the MDH dataset (Supplementary Fig. 21). Thermophilic sequences were oversampled to match the number of mesophilic sequences. Training continued until sequence identity reconverged to the 80% target while the predicted thermal shift stabilised.

For design with the fine-tuned model, the three wild-type queries used for zero-shot design (M7) were supplemented with two additional sequences from MGnify^75^. Variant generation, substitution filtering and pET ranking followed M7. The expanded experimental set comprised 39 designs across the five wild-type backgrounds.

### M10 Molecular Dynamics Simulation and Analysis

Molecular dynamics (MD) simulations were conducted for Lineage 1,2,3 wildtype metagenomic sequences and the 19 finetuned THOR variants with an experimental melting temperature at least 5 °C above their respective wildtype, at and 400 K. Structural models for both the mutated variants and their corresponding wildtype sequences were generated using AlphaFold2^76^ with the full Big Fantastic Database (BFD)^76^ and all five CASP14 models. The structures with the highest average pLDDT score were selected for MD simulations.

Simulations were performed using the GROMACS 2022^77–79^ package with the AMBER99*-ILDN^80^ force field and TIP3P water model^81^. Each protein was placed at the center of a dodecahedron box with a 1 nm boundary to the box edges, solvated, and neutralized with ions. The energy of the solvated systems was minimized using the steepest descent algorithm (50,000 steps, emtol 0.1)^82^ and the Parrinello-Rahman barostat (ref_p = 1 bar, tau_p = 2 ps, compressibility = 4.5 × 10⁻⁵ bar⁻¹)^83^. Periodic boundary conditions were applied in all dimensions for all steps.

The simulations utilized a leap-frog integrator^84^ with a time step of 2 fs, and covalent bonds involving hydrogens were constrained using the LINCS algorithm (lincs_iter = 1, lincs_order = 4)^85^. Non-bonded interactions were handled with a 1 nm cutoff for short-range interactions. Van der Waals interactions employed a Verlet-cutoff scheme (ns_type = grid, nstlist = 10 steps, DispCorr = EnerPres), while electrostatic interactions were calculated using the Particle-Mesh-Ewald (PME) method (pme_order = 4, fourierspacing = 0.16 nm)^86^.

Residue-level root mean square fluctuation (RMSF) profiles were averaged across four independent simulation replicates, and changes relative to the corresponding wild type (WT) were calculated as *ΔRMSF = RMSF_variant_ − RMSF_WT_*. These differences were displayed as residue-by-variant heatmaps with mutation positions annotated. Spatial patterns were characterized using within-profile z-scores of ΔRMSF. Within-lineage heterogeneity was assessed for Lineage 2 using pairwise Pearson correlations between variant ΔRMSF profiles and the occurrence of residues with opposite z-score extremes in different variants. Lineage comparisons used the percentages of variant–residue observations with z > 2 or z < −2. These scores describe deviations from each profile’s mean and were not interpreted as residue-level significance tests.

Hydrogen-bond, salt-bridge and residue-contact counts were averaged within each replicate before comparing variants with their corresponding WT using two-sided Welch t-tests. WT-relative count differences were also summarized within lineages and across all variants using exploratory one-sample t-tests and binomial sign tests; pooled differences were additionally assessed using a Wilcoxon signed-rank test, and differences among lineages using a Kruskal–Wallis test. These count-analysis P values were unadjusted. Pair-level hydrogen-bond and salt-bridge occupancies were compared using two-sided Welch t-tests on replicate occupancies, with Benjamini–Hochberg false discovery rate correction separately for each variant and interaction type (FDR = 0.05).

For structural interpretation, WT-relative interaction occupancy changes were assigned to their residue endpoints and summarized into hydrogen-bond gain, hydrogen-bond loss, salt-bridge gain and salt-bridge loss channels. Each channel was standardized within its lineage, and positions were assigned to their strongest channel when its z-score was ≥2. RMSF and interaction signals were visualized on structural models using UCSF ChimeraX^87^. Recurrent salt bridges with significant occupancy increases in at least two variants were prioritized for schematic lineage-level composite views. Their occupancy increases were summarized over significant supporting variants, with minimum–maximum ranges and the number of supporting variants indicated. Detailed aggregation and structural-selection procedures are provided in the Supplementary Methods.

## Code availability

The source code for THOR is publicly available on GitHub (https://github.com/Umbriona/THOR), and pre-trained model weights are deposited on Hugging Face (https://huggingface.co/Umbriona/THOR). To facilitate reproducibility and accessibility, we provide an executable Google Colab notebook (https://colab.research.google.com/drive/1noVdf3ifoMZVk_RWrobONUiMg3d2DpYm) that enables users to generate thermostability-optimised protein variants directly from input amino acid sequences without local installation. All scripts used for data analysis and generation of the main and supplementary figures are available on GitHub (https://github.com/Umbriona/THOR_Analysis), with the accompanying datasets deposited in Zenodo (doi:10.5281/zenodo.23557423).

