## Supplementary Information for "Cross-family learning transfers thermal adaptation between enzymes"

### Supplementary Expanded methods

#### Phylogenetic trees and sequence sets

Ten COG families were analysed. For each family, we built an alignment comprising natural mesophilic wild types, natural thermophiles, one designed variant per wild type, and outgroup sequences using MAFFT-DASH v7.525[1], trimmed with ClipKIT v2.11.4[2], and inferred a maximum-likelihood tree with IQ-TREE v2.4.0[3]. Across the ten trees, tips comprised 14,637 natural mesophiles, 773 natural thermophiles, 14,619 designed variants, 201 outgroups and 5 test sequences (30,235 tips total). We assigned natural sequences to classes by experimental source-organism growth temperature: mesophiles below 60 °C and thermophiles at or above 60 °C; class labels are complete for every tip. Tip annotations were taken from the pre-clustering sequence metadata table, which covers 100% of tips in all ten trees. The per-tree input metadata was not used, as it was regenerated after two families (COG1012, COG1028) were re-clustered at lower identity thresholds and describes only 35% and 28% of the tips present in those trees.

#### Temperature prediction and predictor calibration

Optimal growth temperature was predicted for every sequence with THOR. The model accepts a maximum input length of 512 residues. Wild types and designed variants were filtered to this ceiling upstream, but natural thermophiles were not: 94 of 773 (12.2%) exceed it and carry no prediction. These sequences are significantly cooler than the remainder (median 65 versus 70 °C, Mann–Whitney $P = 3 \times 10^{-4}$), so the missingness is informative. They were retained in the phylogeny and in clade thermophile status, which derives from the complete class label, but were excluded from any analysis using predicted values. Sequences exceeding the ceiling were not truncated, as cropping would have applied a class-specific transformation confounded with the label being predicted. The predicted axis was validated against natural sequences alone. Separation of natural thermophiles from natural mesophiles gave AUC 0.990 pooled (range 0.981–0.996 across families, $n = 14,637$ mesophiles and 679 thermophiles with predictions). The classification threshold $\tau$ was derived from the natural mesophile distribution as the value giving 95% specificity, $\tau = 47.6$ °C, which yields 95.4% sensitivity. A cutoff of 60 °C, used in earlier analyses, gives 99.4% specificity but only 73.0% sensitivity and was retained solely as a robustness check.

#### Disjoint clade partition

Each tree was partitioned into disjoint monophyletic clades by descending the rooted tree and emitting each maximal subtree containing at most 75 natural tips. Trees were rooted on the outgroup where one was retained and at the midpoint otherwise, as two families (COG0062, COG0108) retain none. Designed variants were pruned before partitioning and each was assigned to its wild type's clade. This yielded 444 analysable clades across the nine families containing natural thermophiles (median 56 tips, 27 pairs), of which 147 contain at least one natural thermophile and 297 contain none. Two alternative criteria were examined. A rooting-free edge-cut partition gave equivalent results (adjusted −0.013, equivalent at ±10 points). A conventional maximum-diameter criterion was rejected: bounding patristic diameter isolates rare divergent sequences into singletons, discarding up to 50% of natural thermophiles, and its estimate converges on the other two as thermophile retention increases.

#### Clade-level enrichment test

For each clade, we computed two outcomes: the fraction of variants with predicted temperature at or above $\tau$ ("hot-variant fraction") and the median per-variant temperature gain over wild type ($\Delta T$). The contrast of interest is the difference in each outcome between clades containing a natural thermophile and clades containing none, with clades as the unit of replication and unweighted means, so that clade size does not reintroduce pseudo-replication. The unadjusted difference was tested by permuting which wild type received which design while holding the tree fixed (10,000 permutations), preserving topology, clade sizes and the marginal distribution of predictions while breaking only the association between clade position and predicted design temperature. Confidence intervals were obtained by bootstrapping clades within group (10,000 resamples). Because clades containing natural thermophiles may have thermophilic amino-acid composition throughout, causing the predictor to score all their sequences higher irrespective of design, the analysis was additionally adjusted for the mean predicted temperature of each clade's wild types in a weighted least-squares model including log clade size, weighted by pairs per clade.

#### Equivalence testing

Equivalence was tested directly by two one-sided tests (TOST) against margins fixed in advance: ±10 percentage points for hot-variant fraction and ±5 °C for $\Delta T$. Equivalence at $\alpha = 0.05$ corresponds to the 90% confidence interval falling entirely within the margin. For each family the smallest margin at which equivalence would be reached, $\delta\_min = |\beta| + z_{0.05} \cdot SE$, is also reported, so that families lacking power for the pre-specified margin still report the resolution they achieve.

#### Per-family analysis and combination

Families differ by more than an order of magnitude in the proportion of designs exceeding $\tau$, so tips were never pooled across families. Each family was analysed separately and the per-family results combined by Stouffer's method weighted by the square root of clade count. COG4188 contains no natural thermophile, cannot contribute to the contrast, and is reported separately.

#### Molecular dynamics trajectory analysis

Molecular dynamics (MD) trajectories were analysed using Python v3.13.5, with MDAnalysis [4,5] for trajectory processing and structural measurements, NumPy and pandas for data handling, SciPy v1.16 for statistical analyses, statsmodels for multiple-testing correction, and Matplotlib for visualisation.

The dataset comprised three wild-type (WT) proteins and 19 THOR-designed variants generated through fine-tuning, distributed across three evolutionary lineages (3, 14 and 2 variants in Lineages 1, 2 and 3, respectively). Trajectories were analysed from 100 ns to the end of each simulation, using every 20th frame. Before calculating root-mean-square fluctuations (RMSF), trajectories were aligned to an average structure computed over the analysis window, using Cα atoms for structural fitting. Per-residue RMSF was then calculated from Cα atom positions and reported in Å.

Protein–protein hydrogen bonds were identified using donor atoms N, NE, NE1, NE2, ND1, ND2, NZ, NH1, NH2, OG, OG1, OH and SG, and acceptor atoms O, OXT, OD1, OD2, OE1, OE2, OG, OG1, OH, ND1, NE2, SD and SG. Hydrogen bonds were defined by a donor–acceptor distance of ≤3.5 Å and a D–H···A angle of ≥150°. Salt bridges were identified between the carboxylate oxygen atoms of Asp and Glu (OD*/OE*) and the side-chain nitrogen atoms of Arg and Lys (NH*/NZ), using a distance threshold of 4.5 Å. Residue contacts were defined as pairs of residues containing at least one pair of protein heavy atoms within 4.5 Å, excluding residues separated by fewer than three positions in the primary sequence.

All structural measurements were performed using MDAnalysis. Exported residue-level RMSF values, framewise interaction counts and residue-pair occupancies were compared with the corresponding WT protein from the same lineage. Where multiple simulation replicates were available, each replicate contributed equally to system-level averages, irrespective of the number of analysed frames. Mutation positions were identified using the accompanying sequence metadata.

#### WT-relative RMSF profiles and heatmaps

RMSF was summarised first within each system–replicate–residue combination and then across the four replicates. For each variant v and residue r, the WT-relative difference was calculated as

$$\Delta\text{RMSF}(v,r) = \text{mean\_rep RMSF}(v,r) - \text{mean\_rep RMSF}(\text{WT},r).$$

We matched residues to the corresponding WT by residue number. Replicate variability was summarised using the sample standard deviation. Heatmaps displayed the unstandardized ΔRMSF values, with variants as rows and residues as columns. Negative and positive values indicated decreased and increased mobility relative to WT, respectively; black squares marked substitution positions. The implementation used a common symmetric colour scale from −4 to +4 in the imported RMSF units.

#### Spatial standardisation and within-lineage heterogeneity

Each variant’s ΔRMSF profile was standardised across residues:

$$z(v,r) = [\Delta\text{RMSF}(v,r) - \text{mean\_r } \Delta\text{RMSF}(v,r)] / \text{SD\_r } \Delta\text{RMSF}(v,r),$$

using the population standard deviation (ddof = 0). Profiles with zero or nonfinite standard deviation were assigned zero scores. These scores identify unusually high or low values relative to the mean of the same profile; they are descriptive rather than inferential statistics.

Within-lineage heterogeneity was evaluated for Lineage 2, as configured in the notebook. Pearson correlations were calculated between all distinct pairs of variant ΔRMSF profiles using residue positions represented in every variant. Correlations were displayed as a matrix and summarised by their median, 25th and 75th percentiles, minimum and maximum. A residue was flagged if at least one variant had $|z| > 2$, and was classified as having opposite extremes if one variant had $z > 2$ and another had $z < -2$. The proportion of flagged positions with opposite extremes was calculated using all flagged positions as the denominator. A separate event-based summary measured the fraction of all flagged variant–residue observations occurring at these positions.

#### Lineage-level distribution of RMSF changes

For each lineage, positive- and negative-tail event rates were calculated as the number of variant–residue observations with $z > 2$ or $z < -2$, respectively, divided by the total number of observations at positions shared by all variants in that lineage. This normalisation is equivalent to averaging the per-variant percentages over the common residue set. The ratio of negative- to positive-tail events was also reported. Position-based rates, defined by whether at least one variant exceeded the threshold, were retained as descriptive summaries but were not used for comparisons across lineages with unequal numbers of variants. Mean unstandardized ΔRMSF and the percentage of positions with a negative lineage-mean ΔRMSF were calculated separately to summarise absolute direction relative to WT.

#### Interaction occupancies and pair matching

Pair occupancy was taken from the exported fraction of analysed frames in which an interaction was present. Multiple entries for the same pair within a replicate, where present, were averaged before replicate aggregation. Unreported pairs were assigned zero occupancy when constructing system means and pair-by-replicate test matrices. Pair identity included both residue numbers and amino-acid identities. Hydrogen-bond pairs retained donor-to-acceptor direction, whereas salt-bridge pairs were ordered by residue number. Consequently, substitution of an endpoint defined a different interaction identity from its WT counterpart. Mean WT occupancy was subtracted from mean variant occupancy to obtain Δoccupancy for each pair.

#### Interaction channels underlying structural mapping

Pairwise Δoccupancy values were assigned to both residue endpoints. Positive changes were summed as gains and negative changes were summed as losses within each variant, residue number and residue identity. Losses were converted to positive magnitudes for comparison of four channels: hydrogen-bond gain, hydrogen-bond loss, salt-bridge gain and salt-bridge loss.

Two aggregation schemes were evaluated in the notebook. The per-position summary averaged the available endpoint-score entries by residue number. The existing ChimeraX exports instead averaged entries by residue number and residue identity, so a substituted position could contribute multiple rows. Within each lineage, each channel was z-scored over the resulting entries using ddof = 0; a constant channel received zero scores. Each entry was assigned to the channel with the largest z-score and was considered mapped when that score was ≥2. Exact ties were resolved in the order hydrogen-bond gain, hydrogen-bond loss, salt-bridge gain and salt-bridge loss. Channel percentages were calculated among mapped entries. For the per-position summary, each residue number contributed once to the final count. An exploratory Pearson chi-square test compared the numbers attributed to salt bridges versus hydrogen bonds among the three lineages using a $3 \times 2$ contingency table. These channel scores quantify the spatial concentration of interaction changes and do not measure total interaction abundance or establish residue-level significance.

#### Absolute interaction counts

For hydrogen bonds, salt bridges and residue contacts, framewise counts were averaged separately within each replicate. Each variant's effect was defined as the mean of its four replicate averages minus the mean of its WT's four replicate averages, in units of interactions. Individual variant–WT comparisons used two-sided Welch t-tests with the four independent replicate averages in each group.

Within each lineage, the distribution of variant effects was summarised by its mean, range and number of positive differences. Two-sided one-sample t-tests evaluated a mean effect of zero, and two-sided exact binomial tests evaluated whether the fraction of positive effects differed from 0.5. The same tests were applied to the pooled set of 19 variant effects, with each variant receiving equal weight. The pooled analysis also included a two-sided Wilcoxon signed-rank test against zero. A Kruskal–Wallis test compared effect distributions among lineages. P values from this count-analysis section were not adjusted for multiple testing; comparisons with $P < 0.05$ were tabulated. The supplementary count plot showed individual variant effects and lineage means ± standard error across variants. Because effects within a lineage share an estimated WT baseline, lineage and pooled tests were treated as exploratory summaries conditional on that baseline.

#### Identification of recurrent interaction gains

Hydrogen-bond and salt-bridge occupancies were tested separately. For each lineage and interaction type, the union of pair identities observed in WT or any variant defined the testing set. Two-sided Welch t-tests compared the four variant replicate occupancies with the four WT replicate occupancies. Benjamini–Hochberg correction was applied to valid P values across pairs separately for each variant and interaction type, at FDR = 0.05. A significant gain required rejection after correction and a positive mean Δoccupancy. Identical constant samples were assigned $P = 1$; other nonfinite test results were excluded from correction. Recurrence was summarized as the number of variants with significant gains and that number divided by the number of variants with a finite test for the pair. This was a recurrence summary, rather than an additional lineage-level hypothesis test.

For WT interaction-focus views and representative variant close-ups, pairs were eligible when significant gains occurred in at least 50% of tested variants or in at least two variants, and significant gains outnumbered significant losses. Eligible pairs were ranked by significant-gain fraction, significant-gain count and mean occupancy increase among variants with positive changes, with pair label used to break ties. Up to three pairs per lineage and interaction type were exported for WT focus views, and up to two for representative variant close-ups. Among significant supporting variants with an available structure, the representative was selected by decreasing Δoccupancy, decreasing variant occupancy and increasing adjusted P value, with variant identifier as the final tie-breaker.

#### Structural visualization

WT structures were oriented relative to the Lineage 1 WT using least-squares Cα superposition of atoms matched by chain identifier, residue number and insertion code. Representative variant structures were similarly oriented to the configured Lineage 1 reference. Structural figures were prepared using UCSF ChimeraX[6]. The signed RMSF export averaged raw ΔRMSF over available variant entries grouped by residue number and residue identity, then standardized these averages within each lineage. Its configured color breaks were $|z| = 1$, 2 and 3, with entries below $|z| = 1$ neutral. The combined interaction display used the dominant-channel scheme described above, with entries below $z = 2$ neutral. The saved RMSF and interaction exports therefore retain the residue-identity grouping described above.

Interaction close-ups displayed the two endpoint residues as sticks against a neutral cartoon. Substituted endpoints were orange and unchanged endpoints blue. Dashed connectors in representative static structures joined the closest available heavy side-chain atoms, with Cα used as a fallback where side-chain atoms were absent. These connectors illustrated the selected residue pair and were not used to determine trajectory interaction occupancy.

#### Recurrent salt-bridge composites and quantitative occupancy plot

Composite views retained salt bridges with FDR-significant positive occupancy changes in at least two variants. No additional minimum recurrence fraction or mean occupancy increase was imposed. Pairs were ranked by significant-gain fraction, significant-gain count and mean positive Δoccupancy. Up to four pairs per lineage were retained, proceeding through this ranking and skipping pairs that required incompatible amino-acid identities at an already assigned schematic position.

For each significant supporting variant, a pair was classified as directly introduced if at least one endpoint was substituted, or native/conformational if neither endpoint was substituted. A pair was classified as mixed if its supporters included both categories. These are sequence-annotation categories, rather than independent demonstrations of a physical mechanism. The aligned WT provided the common schematic backbone, and introduced endpoint identities were represented using ChimeraX side-chain substitutions. Composite views could combine substitutions from different variants and therefore did not represent a sampled or energy-minimised multimutant structure.

The companion occupancy plot used the same selected pairs. Each point showed the mean Δoccupancy among variants with FDR-significant gains, multiplied by 100 to express percentage-point changes. Horizontal lines extended from the minimum to the maximum Δoccupancy among those variants; these were ranges, not confidence intervals. Marker size encoded the number of significant supporters, color identified lineage, and shape distinguished directly introduced, native/conformational and mixed pairs. Labels reported significant supporters divided by the number of tested variants. These conditional means describe the significant supporting subset rather than all variants in a lineage.

### Supplementary figures

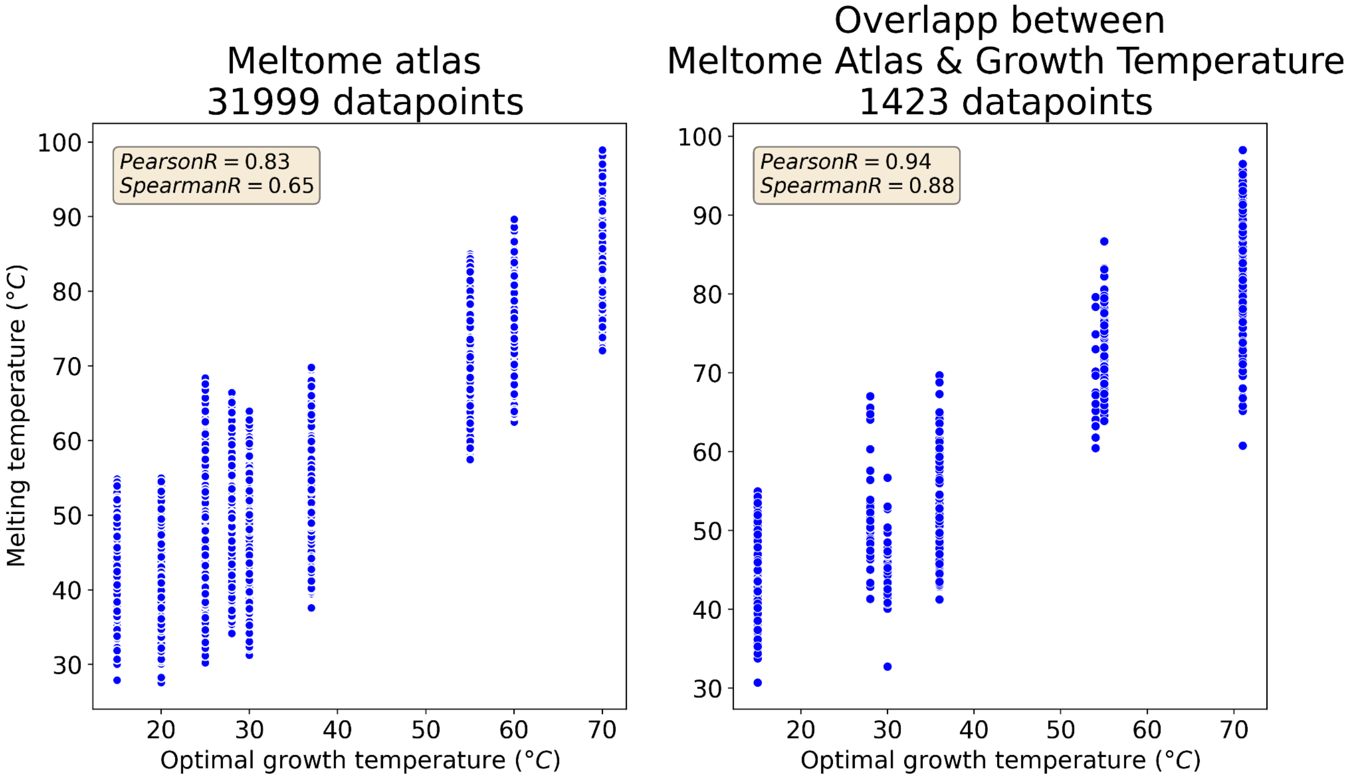


**Supplementary Figure 1.** Correlation between Melting temperature and optimal growth temperature. The first plot shows the data from the Meltome atlas, and the second plot shows the overlapping sequences between the Meltom atlas and the growth temperature data set.

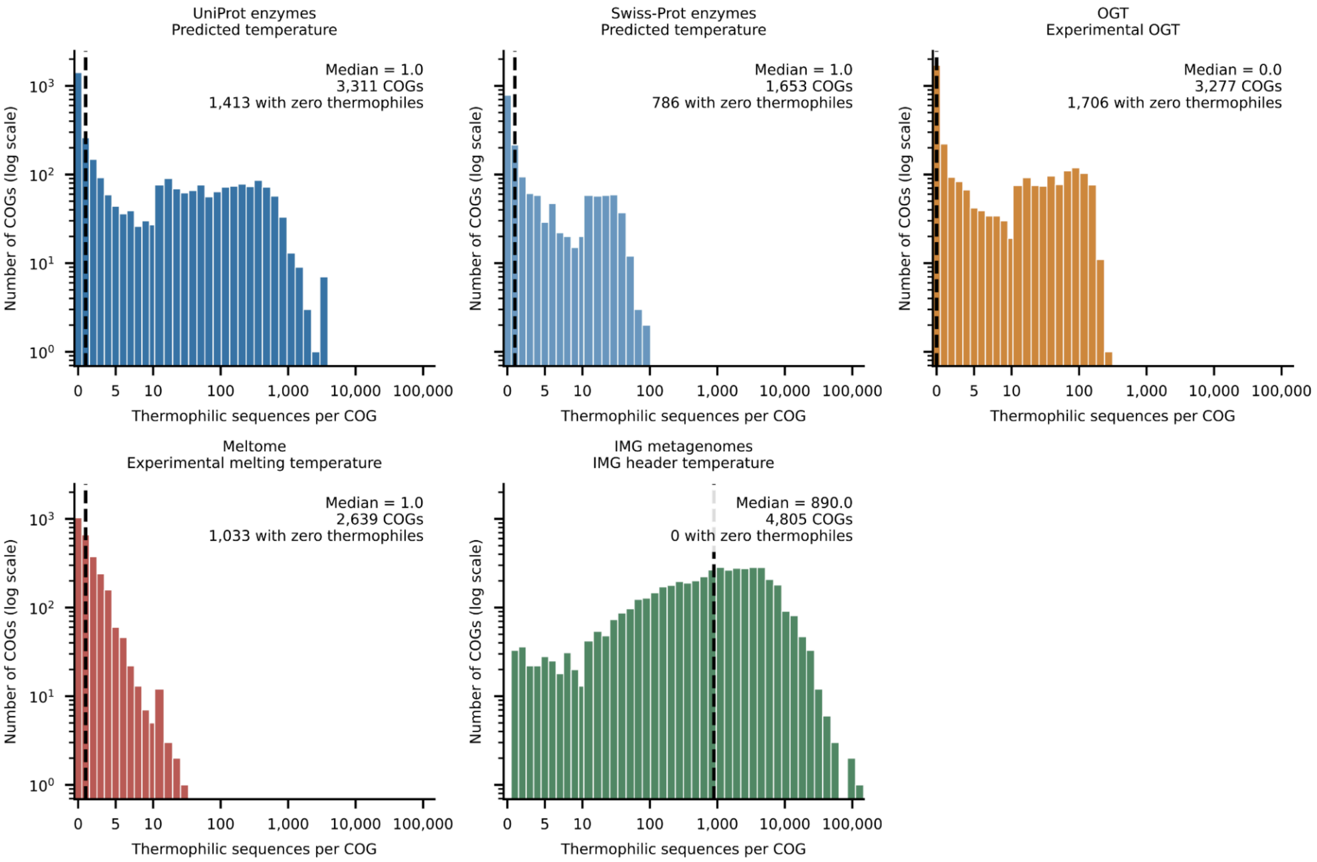


**Supplementary Fig 2.** Distribution of thermophilic sequence counts across COGs. Histograms show the number of thermophilic protein sequences per root-level cluster of orthologous groups (COG) in the UniProt enzyme, Swiss-Prot enzyme, OGT, Meltome, and IMG metagenomic datasets. Sequences were classified using a threshold of ≥60°C, applied to predicted temperatures for UniProt and Swiss-Prot, experimental optimal growth temperatures for OGT, experimental melting temperatures for Meltome, and temperatures recorded in the IMG FASTA headers. The Meltome classification therefore reflects protein melting temperature rather than organismal growth temperature. COG assignments were obtained from EggNOG annotations, counting each sequence ID once per dataset and excluding ambiguous root assignments. Swiss-Prot annotations were additionally restricted to entries with an EC assignment. Each histogram includes all eligible COGs represented in its dataset, including those with zero thermophilic sequences. Dashed lines indicate median counts: UniProt, 1; Swiss-Prot, 1; OGT, 0; Meltome, 1; and IMG, 890. The x-axis is linear near zero and logarithmic above 10, with wider bins at larger counts; the y-axis shows the number of COGs per bin on a logarithmic scale.

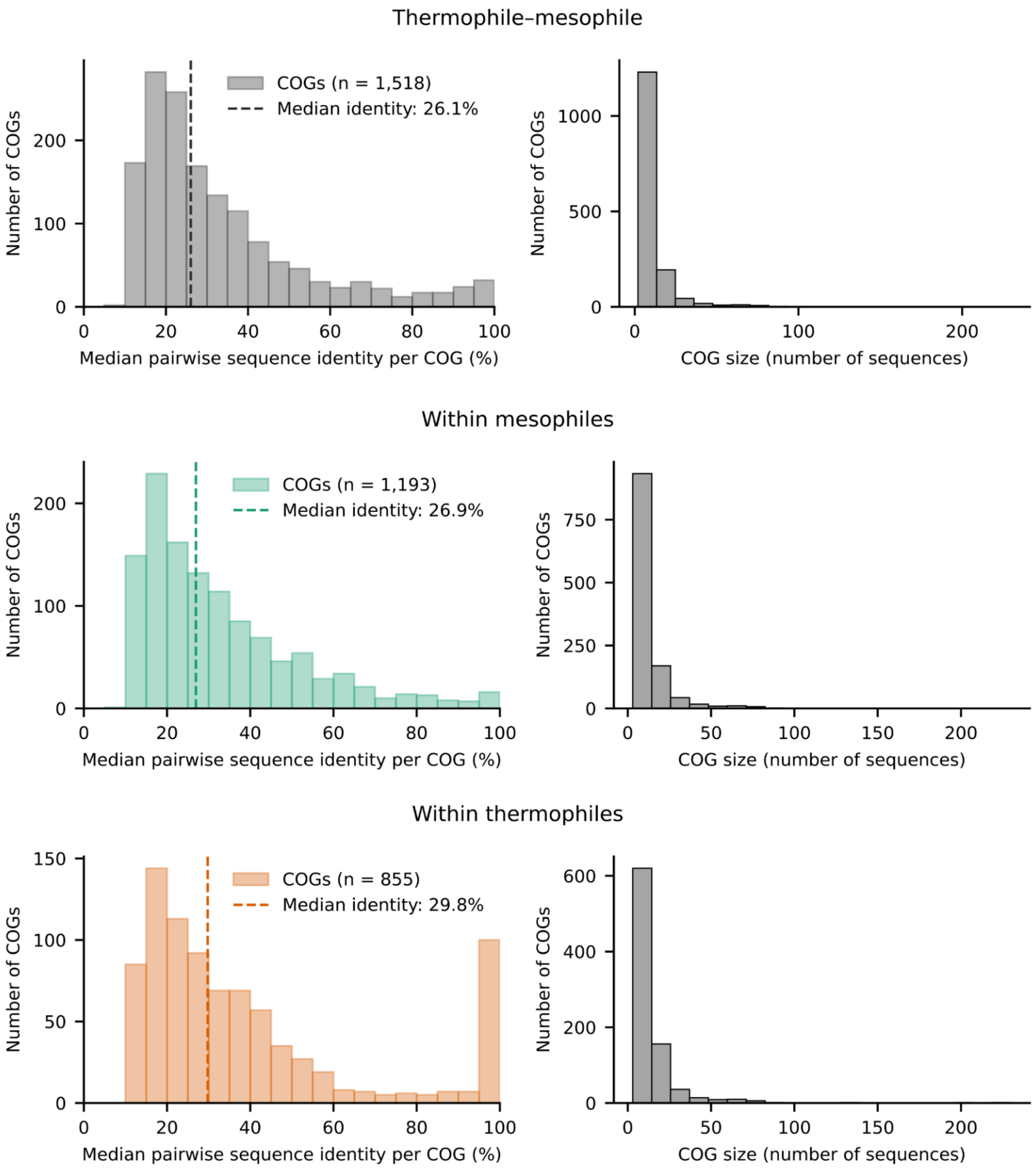


**Supplementary Figure 3.** Sequence identity within orthologous groups across temperature classes. Distributions of pairwise sequence identity between thermophilic and mesophilic proteins (≥60 °C and <60 °C, respectively), within thermophiles, and within mesophiles from the Meltome Atlas. For each COG, sequences were aligned using Clustal Omega and the median pairwise identity was calculated, excluding self-comparisons and duplicate symmetric comparisons. In each figure, the left panel shows the distribution of median identity across COGs; the dashed line indicates the median across COGs. The right panel shows the corresponding distribution of COG sizes.

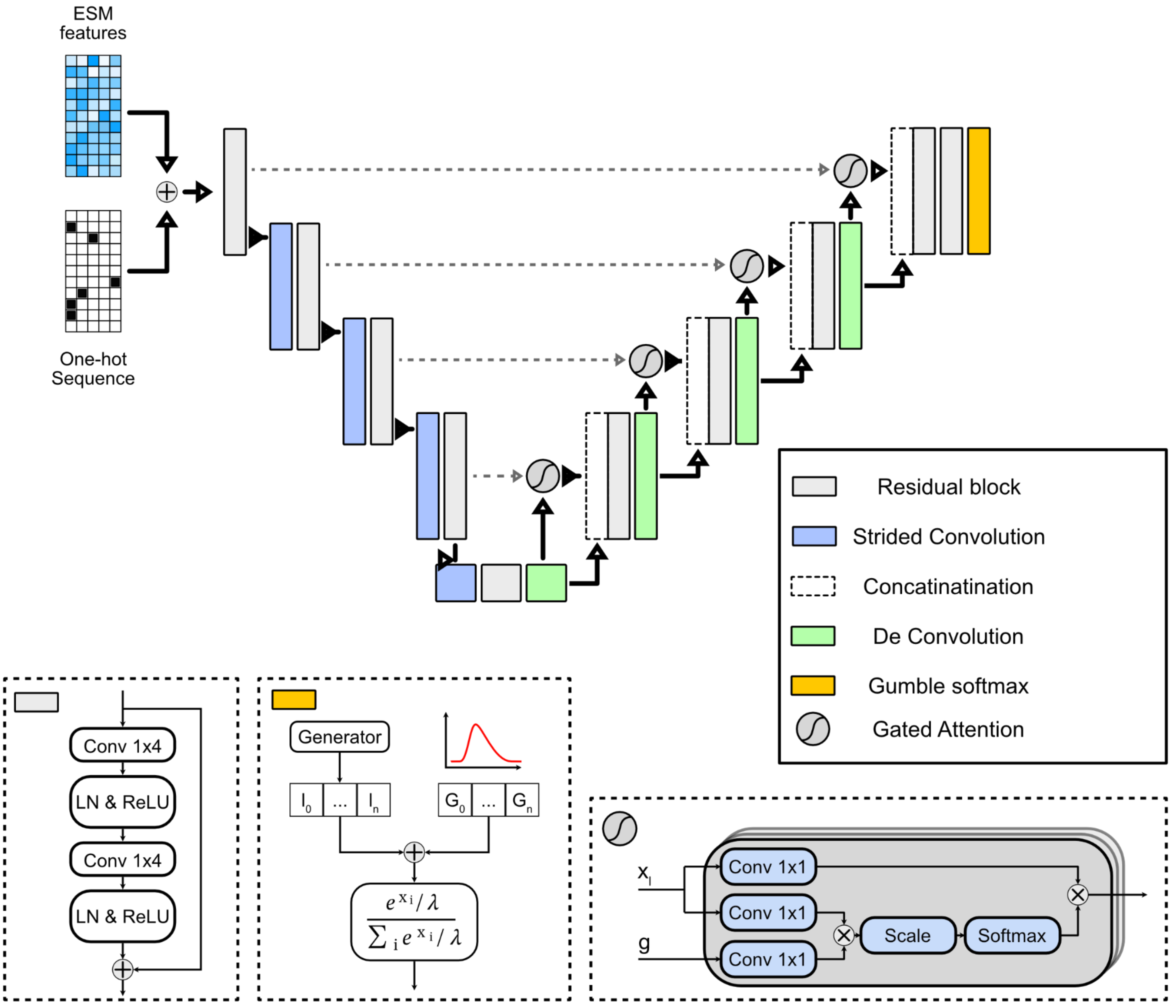


**Supplementary Figure 4.** Architecture of the Gated Attention U-Net generator for protein sequence generation. The network takes as input a one-hot encoded amino acid sequence and ESM-derived likelihood features (blue and black grids on the left). These are concatenated and passed through an encoder-decoder U-Net structure with skip connections. The encoder (blue blocks) progressively reduces spatial resolution, extracting hierarchical features, while the decoder (green blocks) reconstructs sequence representations. Gated attention units (gray circles) modulate skip connections by learning relevance weights, improving information flow between encoder and decoder stages. At the bottleneck, the network captures global contextual information. The output layer applies a convolution followed by Gumbel-Softmax activation to produce a differentiable approximation of categorical distributions over amino acids, enabling sequence generation while maintaining end-to-end differentiability. Additional modules include 1×1 convolutions and scaling operations for attention weighting.

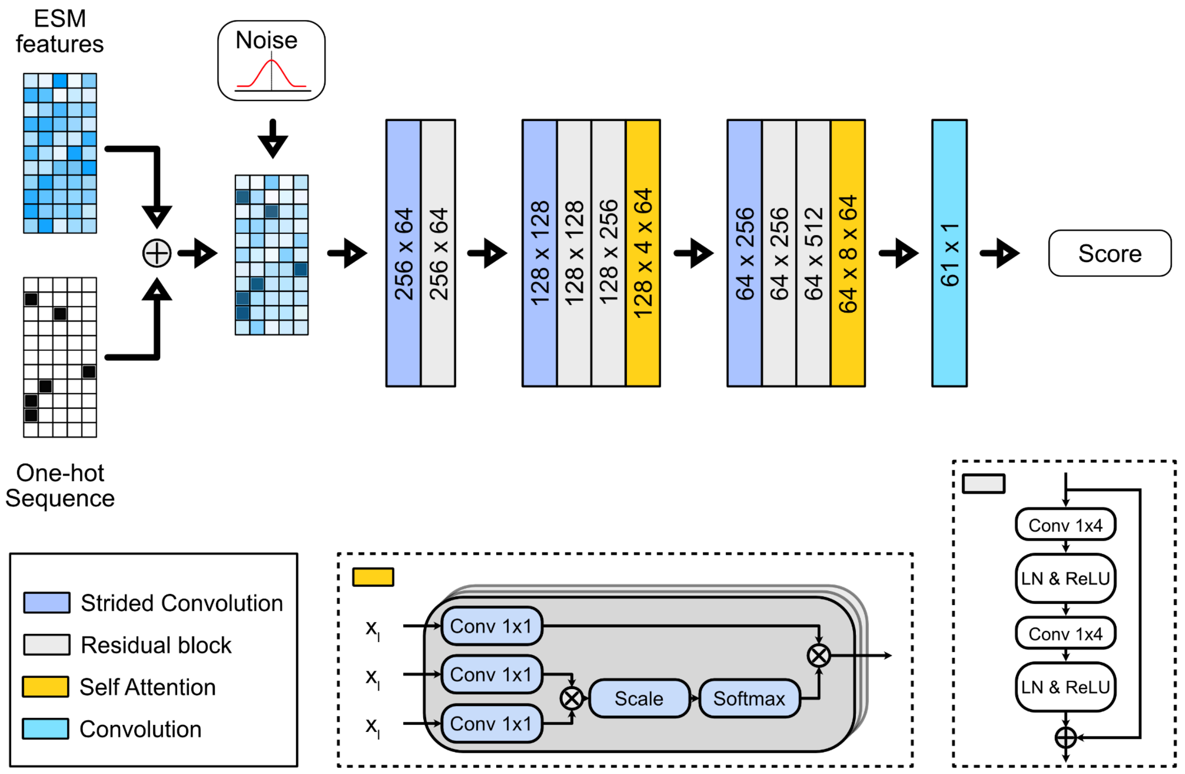


**Supplementary Figure 5.** Architecture of the discriminator network for protein sequence evaluation. The input consists of a one-hot encoded amino acid sequence and ESM-derived likelihood features, which are added together and perturbed with Gaussian noise for regularization. The network processes this input through a series of strided convolutional layers (blue blocks) to reduce dimensionality, followed by residual blocks (gray blocks). Self-attention layers (yellow blocks) are interleaved to capture long-range dependencies across sequence positions. The final convolutional layer (cyan block) outputs a per-position score, The final convolutional layer (cyan block) outputs a sequence of local realism scores (PatchGAN-style), where each score corresponds to a specific region of the input sequence. Inset diagrams illustrate the structure of self-attention and residual blocks used within the architecture.

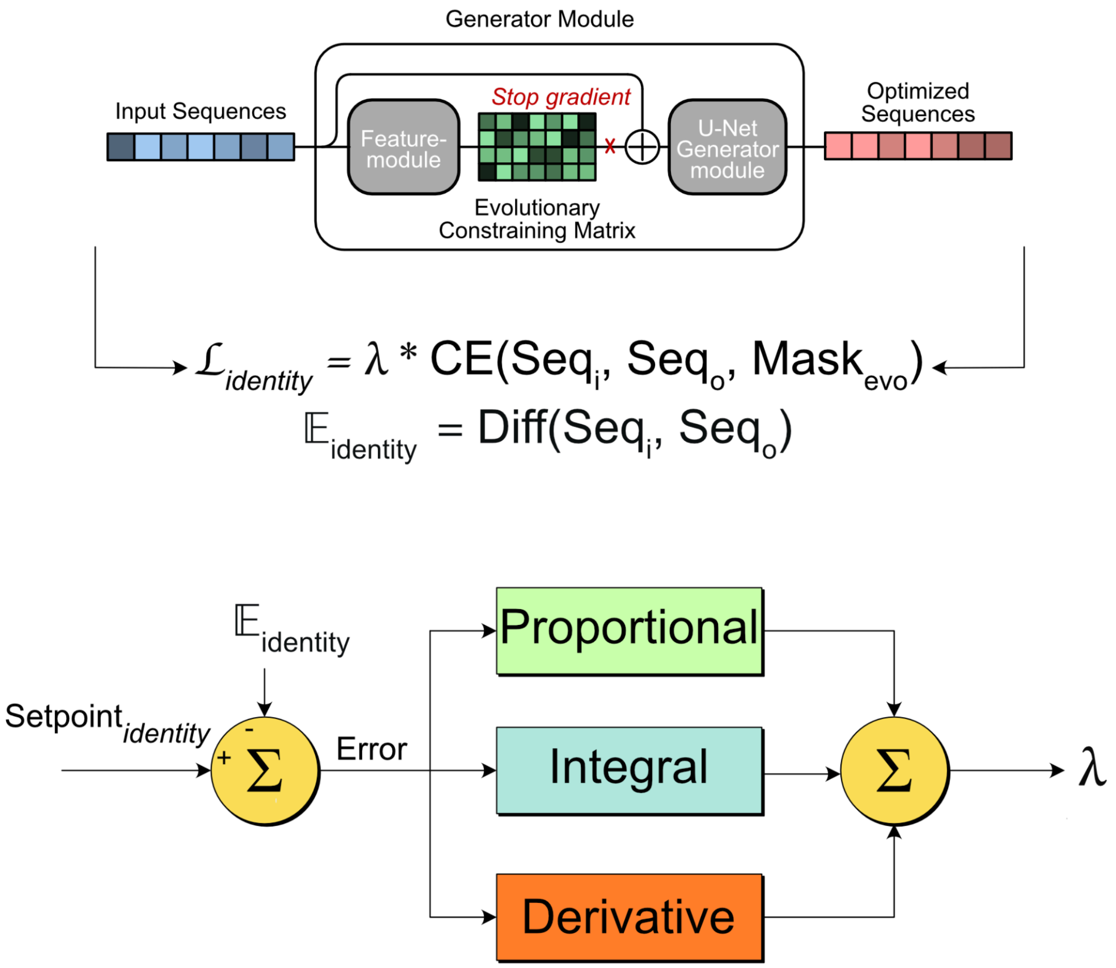


**Supplementary Figure 6.** PID-controlled identity loss used to enforce a target sequence identity during generator training. A proportional–integral–derivative (PID) controller was used to regulate the sequence identity between the generated optimized sequences and their corresponding input (wild-type) sequences. For each training step, the observed sequence identity, $E_{identity}$, was calculated from the difference between the input sequence ($Seq_0$) and the generated sequence ($Seq_i$) and compared with a predefined target identity (setpoint). The resulting error was processed by the proportional, integral, and derivative components of the PID controller to determine λ, which controls the relative contribution of the identity loss to the generator training objective. The identity loss was calculated as a masked cross-entropy loss between the generated and input sequences. The evolutionary mask ($Mask_{evo}$) was derived from the evolutionary constraining matrix produced by the feature module. The constraining matrix was combined element-wise with the input sequence and reduced across the amino-acid dimension to generate a position-specific masking vector, thereby restricting the identity loss to evolutionarily constrained sequence positions. Gradients were stopped through the evolutionary constraining matrix when calculating this loss. By dynamically adjusting λ, the PID controller provides feedback control over the degree of sequence modification and maintains the generated sequences near the desired target identity during training.

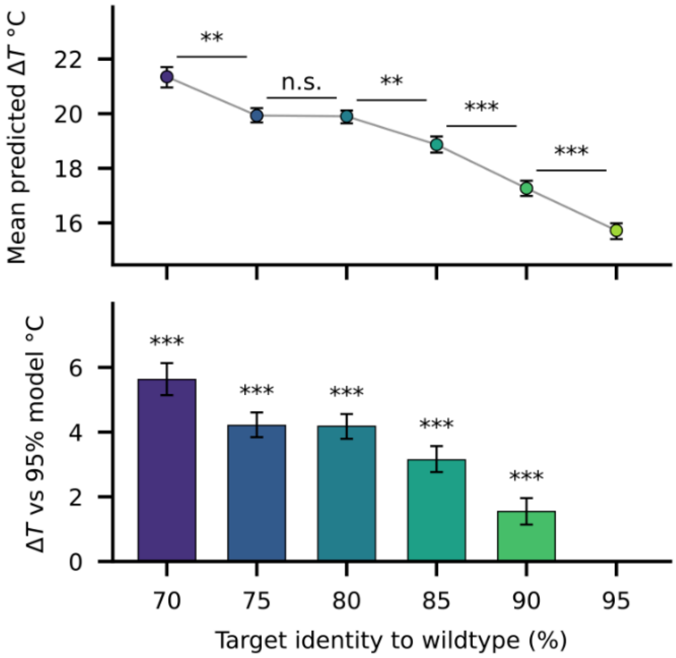


**Supplementary Figure 7.** The predicted thermal shift saturates below ~80 % target identity to the wildtype.(a) Mean predicted ΔT per target identity. Error bars are 95 % confidence intervals from a moving-block bootstrap, which preserves the autocorrelation between successive evaluation batches. Brackets give two-sided Wilcoxon rank-sum tests between neighbouring identity targets, computed on batches thinned to their decorrelation lag and Holm-corrected across the five comparisons (** $q < 0.01$, *** $q < 0.001$, n.s. not significant). Predicted ΔT falls monotonically as the identity target is tightened, from 21.4 °C at 70 % to 15.7 °C at 95 % identity, but is indistinguishable between the 75 % and 80 % models (difference 0.03 °C, 95 % CI −0.31 to +0.42 °C, $P = 0.43$), whereas every other 5-point step changes ΔT by 1.0–1.6 °C ($q \leq 0.009$).

(b) Predicted ΔT of each model relative to the 95 % identity model, with the pairwise moving-block bootstrap CI and Holm-corrected rank-sum q-value. Relaxing the target from 95 % to 80 % identity buys 4.2 °C (95 % CI 3.8–4.6 °C); relaxing it further to 75 % buys nothing measurable, and to 70 % a further 1.4 °C. Even the 95 % identity model, which changes on average fewer than one residue in twenty, produces sequences with a mean predicted increase above 15 °C.

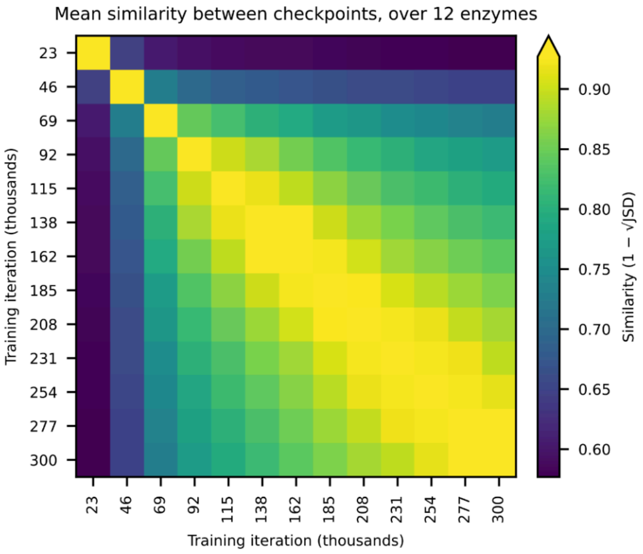


**Supplementary Figure 8.** THOR's per-residue predictions. Mean pairwise similarity between the model's output at 13 training checkpoints, taken every three epochs (23,100 iterations) across the run. At each checkpoint, the model scored the same 12 PET hydrolases (which it had never seen), producing a per-residue probability distribution over the 21 amino-acid classes for each enzyme. Similarity between two checkpoints is $1 - \sqrt{JSD}$, where JSD is the Jensen–Shannon divergence between the two distributions averaged across residue positions; the square root is taken because it, unlike JSD itself, is a metric. Values are averaged over the 12 enzymes, and 1 denotes identical predictions. The diagonal equals 1 by construction; the colour scale spans off-diagonal values only.

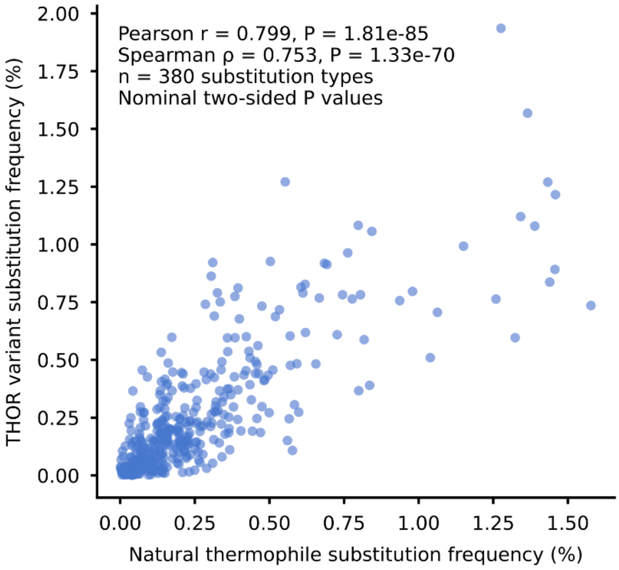


**Supplementary Figure 9.** Correlation between natural thermophile and THOR substitution frequencies. Each point represents one of 380 directed amino acid substitutions, comparing pooled mesophile-to-thermophile frequencies with mesophile-to-THOR frequencies. Unchanged residues were excluded; zero-frequency substitutions were retained. Frequencies were positively correlated (Pearson $r$ = 0.799, $P$ = 1.81 × $10^{-85}$; Spearman $\rho$ = 0.753, $P$ = 1.33 × $10^{-70}$). Two-sided $P$ values are nominal and do not account for dependence between substitution frequencies.

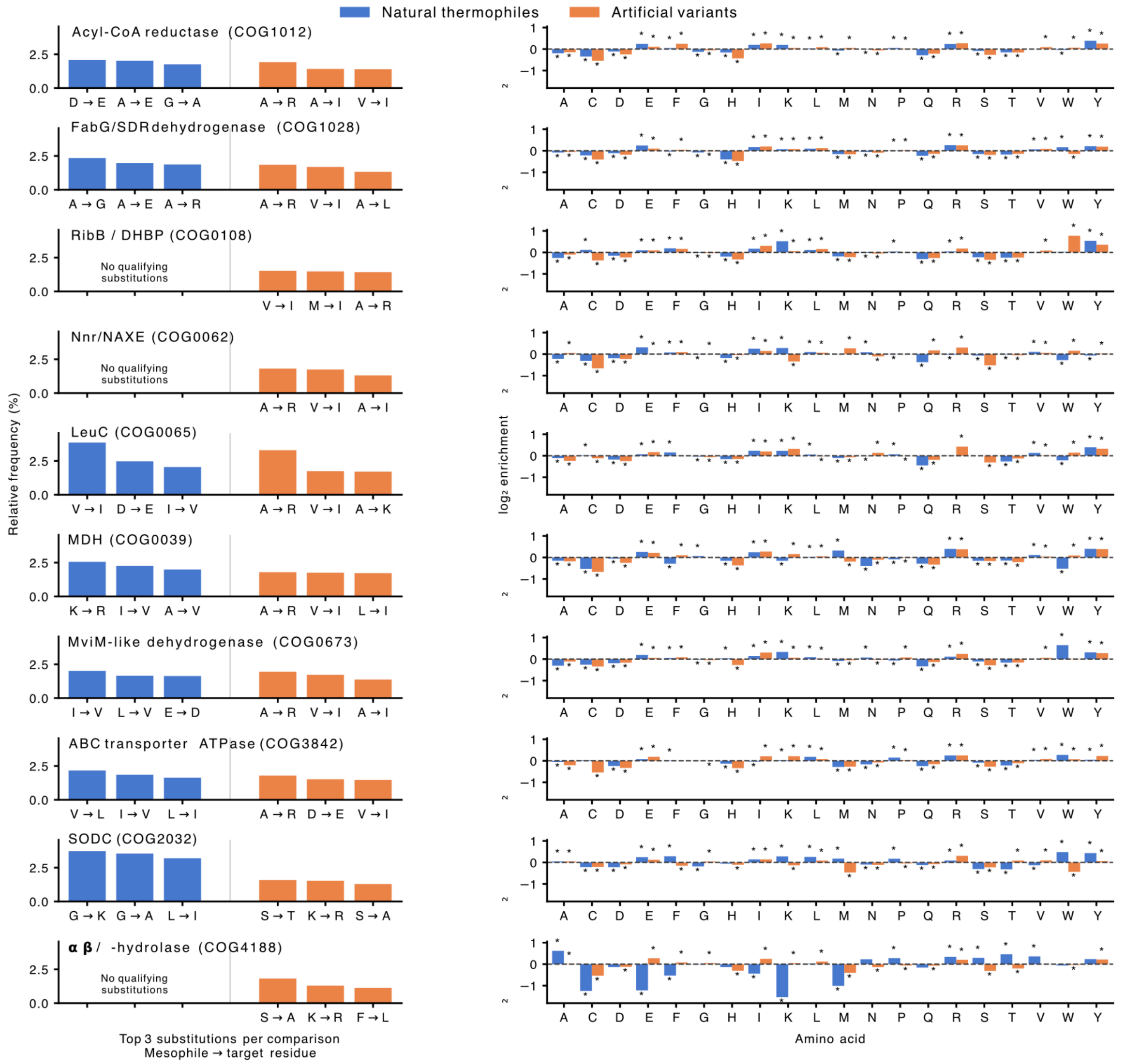


**Supplementary Figure 10.** Family-specific substitutions and amino acid enrichment in natural thermophiles and THOR variants in test set families. Each row represents one protein family. **Left:** Top three mesophile-to-thermophile (blue) and mesophile-to-THOR (orange) substitutions, selected independently and normalized within each family and comparison. Natural substitutions were inferred from temperature-associated alignment positions; THOR substitutions were counted from paired wildtype–variant sequences. "No qualifying substitutions" indicates that no alignment met the group-size and residue-association requirements needed to identify substitutions. **Right:** $Log_2$ amino acid enrichment relative to mesophiles from the same family. Asterisks indicate adjusted $P < 0.05$ from two-sided Fisher's exact tests with Benjamini–Hochberg correction across the 40 comparisons per family.

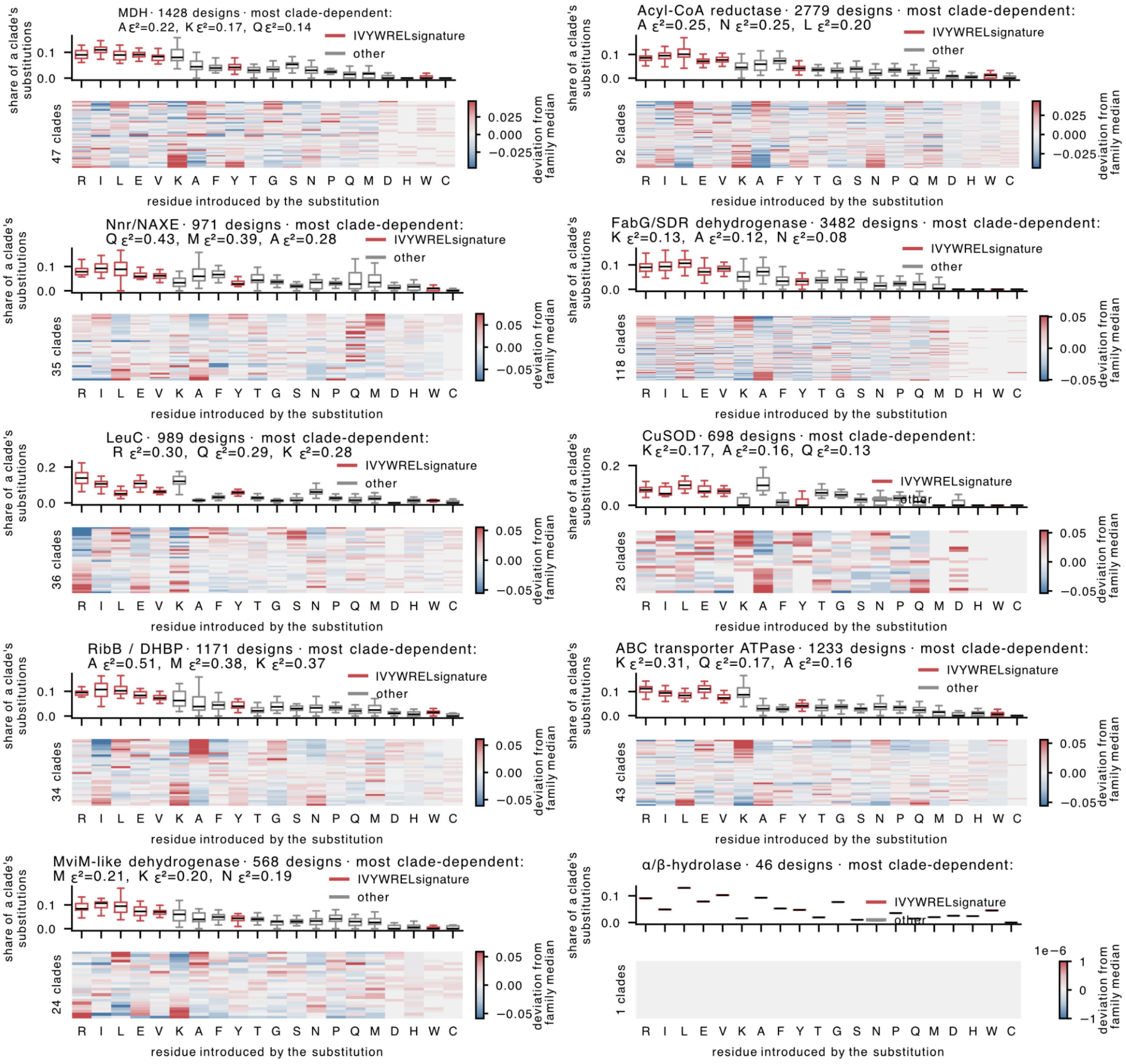


**Supplementary Figure 11.** Designed substitution spectra differ between clades within every family in the test set. Frequencies are conditional on a substitution having occurred. They sum to one across the twenty residues within a design, so substitution rate is divided out. Residues are ordered by pooled frequency, commonest first, identically in every panel. Upper panel: share of a clade's substitutions introducing each residue, one observation per clade (that clade's median); box, median and interquartile range; whiskers, 1.5× IQR; outliers omitted. Red, the seven IVYWREL signature residues; grey, the other thirteen. Lower panel: one row per clade, coloured by that clade's deviation from the family median, symmetric about zero with limits at the 98th percentile of |deviation|. Clades are ordered by the first principal component of the matrix shown; this ordering is derived from the same data and is descriptive only. Clade dependence was tested per family and residue by Kruskal–Wallis, against a null shuffling clade labels within each family (2,000 permutations); 170 of 180 tests reach $q < 0.05$. K, A, Q and M vary most between clades (mean $\varepsilon^2$ 0.18–0.23), V, W and C least (≈0.03).

*α/β-hydrolase (j) holds a single clade, so its boxes collapse to lines and no deviation panel is defined. Per family: 46–3,482 designs across 1–118 clades (13,365 and 453 in total). Axes and colour limits are scaled per panel.

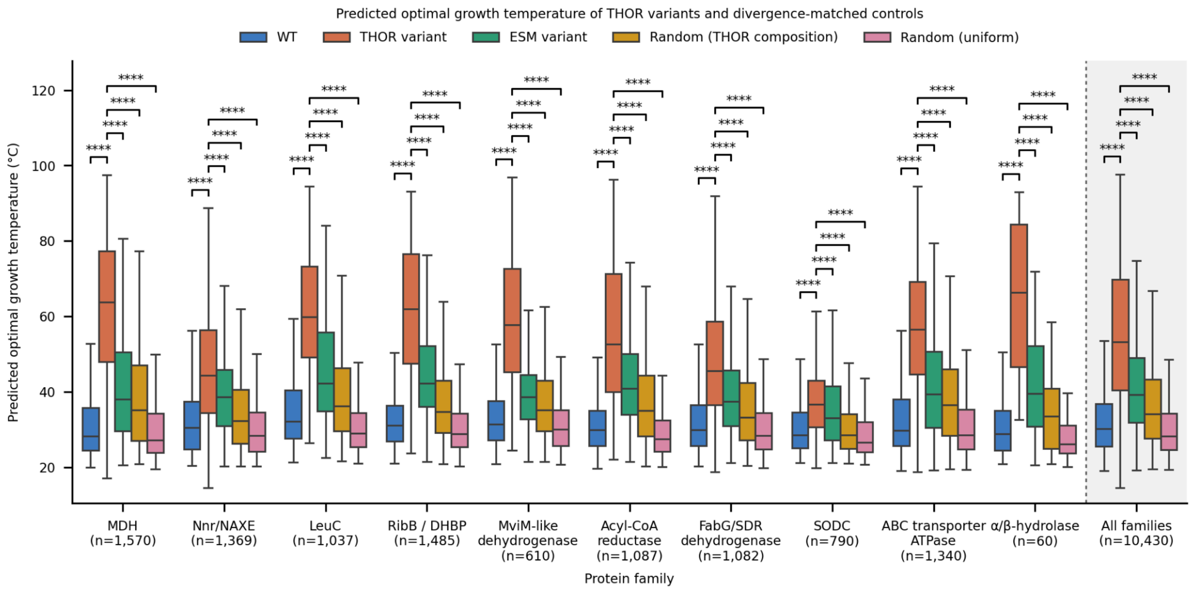


**Supplementary Figure 12.** Predicted optimal growth temperature of THOR variants and divergence-matched controls in the test set, by protein family. Predicted optimal growth temperature (pOGT) for 10,430 mesophilic wildtype enzymes drawn from ten COG families and for four designed variants of each, one per wildtype. Boxes show the median and interquartile range with whiskers at 1.5 × IQR; outliers are omitted. The shaded **All families** panel pools every family's sequences and is a re-analysis of the same data rather than an independent sample. **n** is the number of matched wildtypes in each panel, which is also the number of sequences in each box.

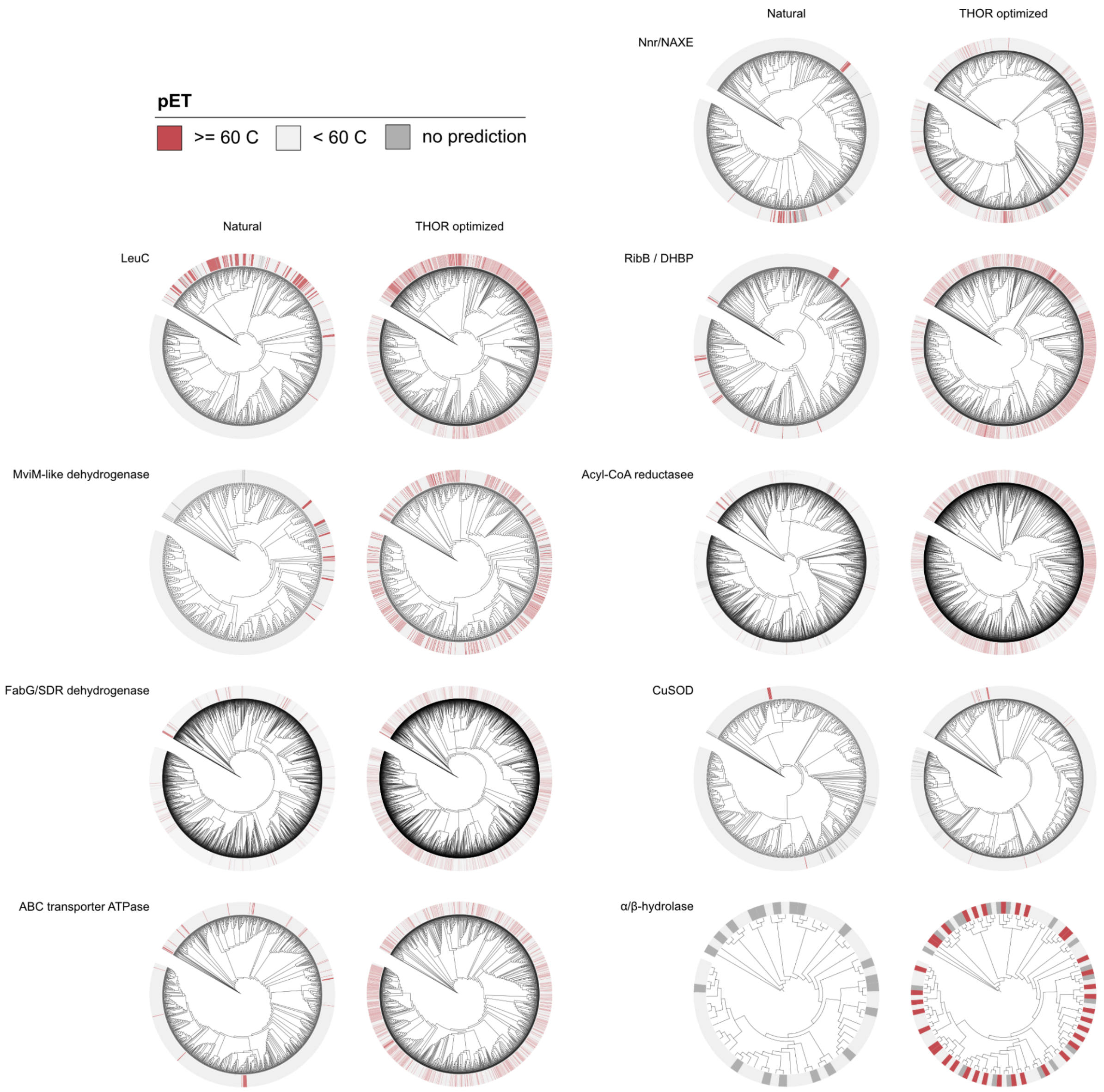


**Supplementary Figure 13.** Phylogenetic trees before and after adding designed variants. Ten COG test set families, five per column; for each, the left tree is built from natural sequences alone and the right from identical inputs plus one designed variant per mesophilic wild type. The pairs share 98.8–100% of their natural sequences, so they differ only by the variants. Outer strip: predicted optimal growth temperature: red ≥ 60 °C, pale < 60 °C, grey no prediction. Grey tips are natural thermophiles exceeding the model's 512-residue limit; they are experimentally thermophilic but unscoreable, not cold. Inner strips give sequence class. Tips predicted ≥ 60 °C rise from 3.7% to 20.7% across all families (COG2032 is the exception, 0.7 → 1.1%). Descriptive only; tests are in Figureures S2 and S3.

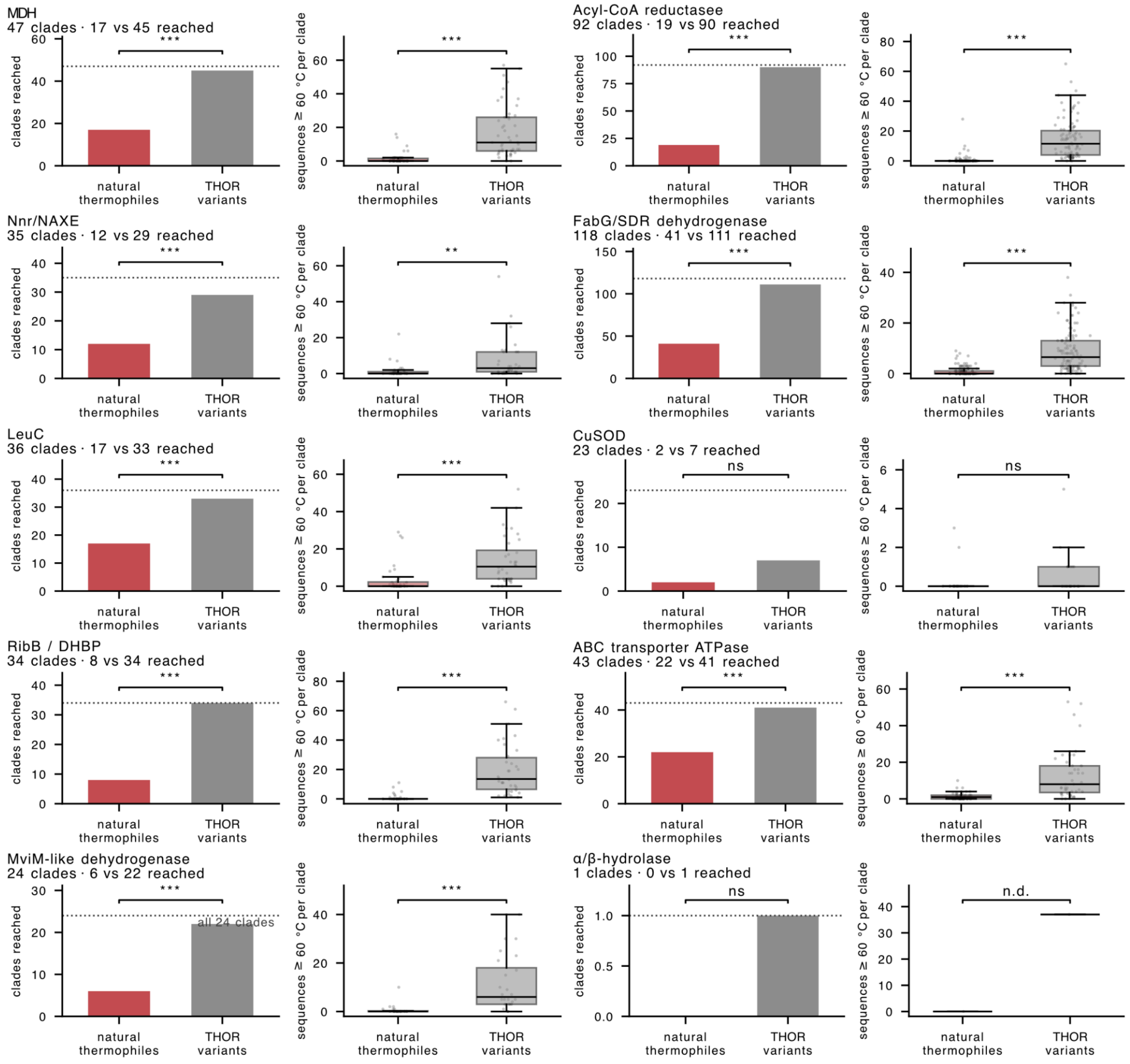


**Supplementary Fig 14.** Natural thermophiles and designed variants as two routes to thermophilicity in test set families. Each tree is partitioned into disjoint monophyletic clades (453 analysed). Left panel of each pair: clades containing ≥ 1 natural thermophile, defined by experimental growth temperature, versus clades containing ≥ 1 variant predicted ≥ 60 °C. Right panel: how many such sequences each route contributes per clade (points, clades; boxes, median and quartiles). Both routes are scored on the same clades, so the comparison is paired and chi-square does not apply. Asterisks give McNemar's exact test (left) and Wilcoxon signed-rank (right): *** $P < 0.001$, ns not significant, n.d. not computable. Pooled, 139 clades are reached by both routes, 274 by designs only and 5 by naturals only ($P = 3 \times 10^{-74}$). At matched sequence counts, designs still reach 246 clades [233, 260] against 144.

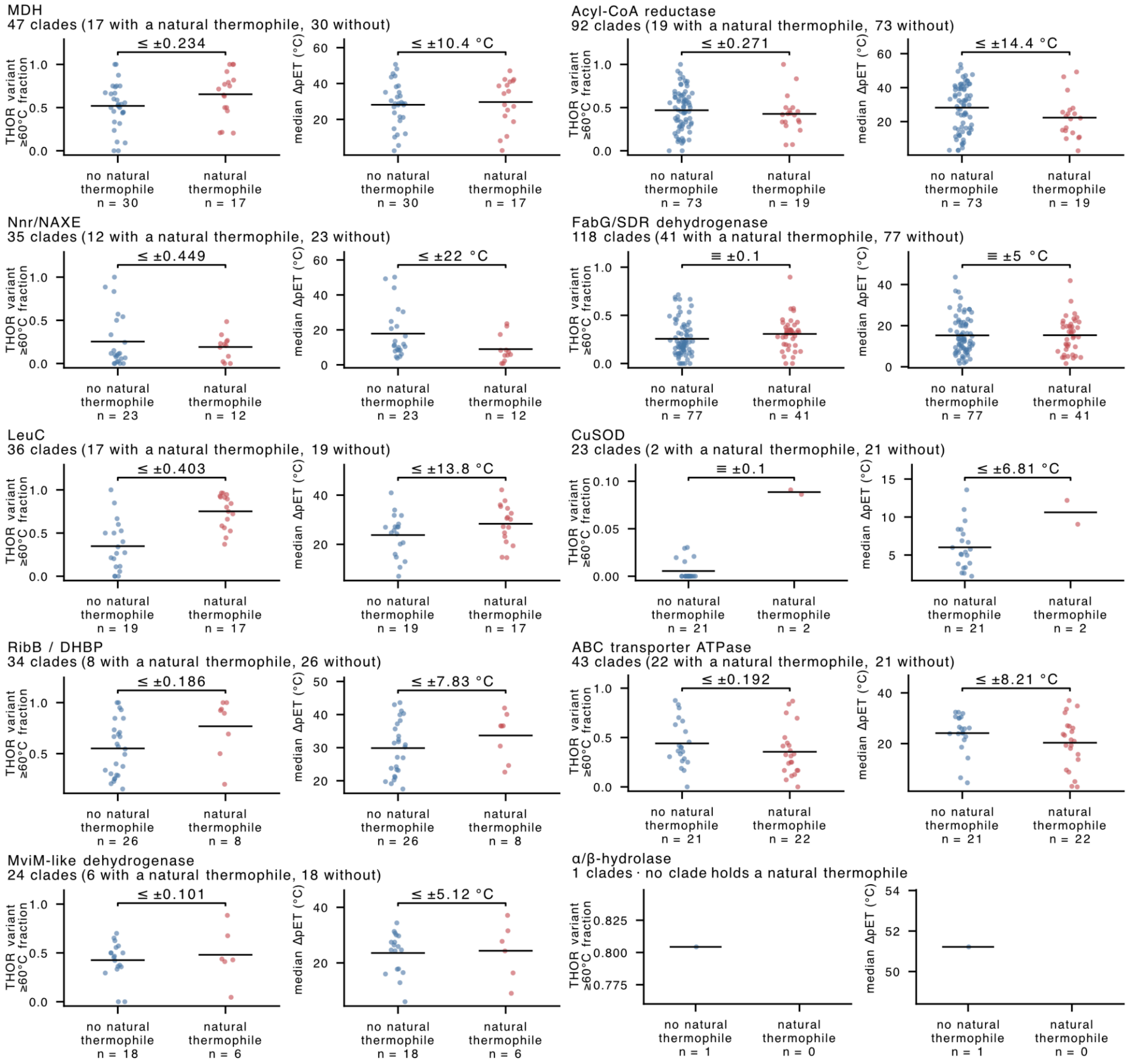


**Supplementary Figure 15.** Designs raise predicted thermophilicity independently of local natural thermophiles in the test set families. Clades are split by whether they contain a natural thermophile, defined experimentally. Left panel of each pair: fraction of a clade's variants predicted ≥ 60 °C. Right panel: median per-variant gain over wild type (ΔT). Points are clades, bars are group means, group sizes beneath the axis (452 clades: 144 with a natural thermophile, 308 without). The bracket gives the difference between groups after adjustment for the mean predicted temperature of the clade's wild types. without which the predictor's tendency to score thermophile-rich neighbourhoods higher produces a difference on its own. ≡ marks equivalence to zero at the stated margin by two one-sided tests; otherwise the bracket gives the smallest margin that family can resolve. Pooled, the adjusted difference is −0.016 (90% CI −0.061 to +0.029), equivalent at ±10 percentage points ($P = 0.001$); unadjusted it is +0.076 ($P = 1 \times 10^{-4}$).

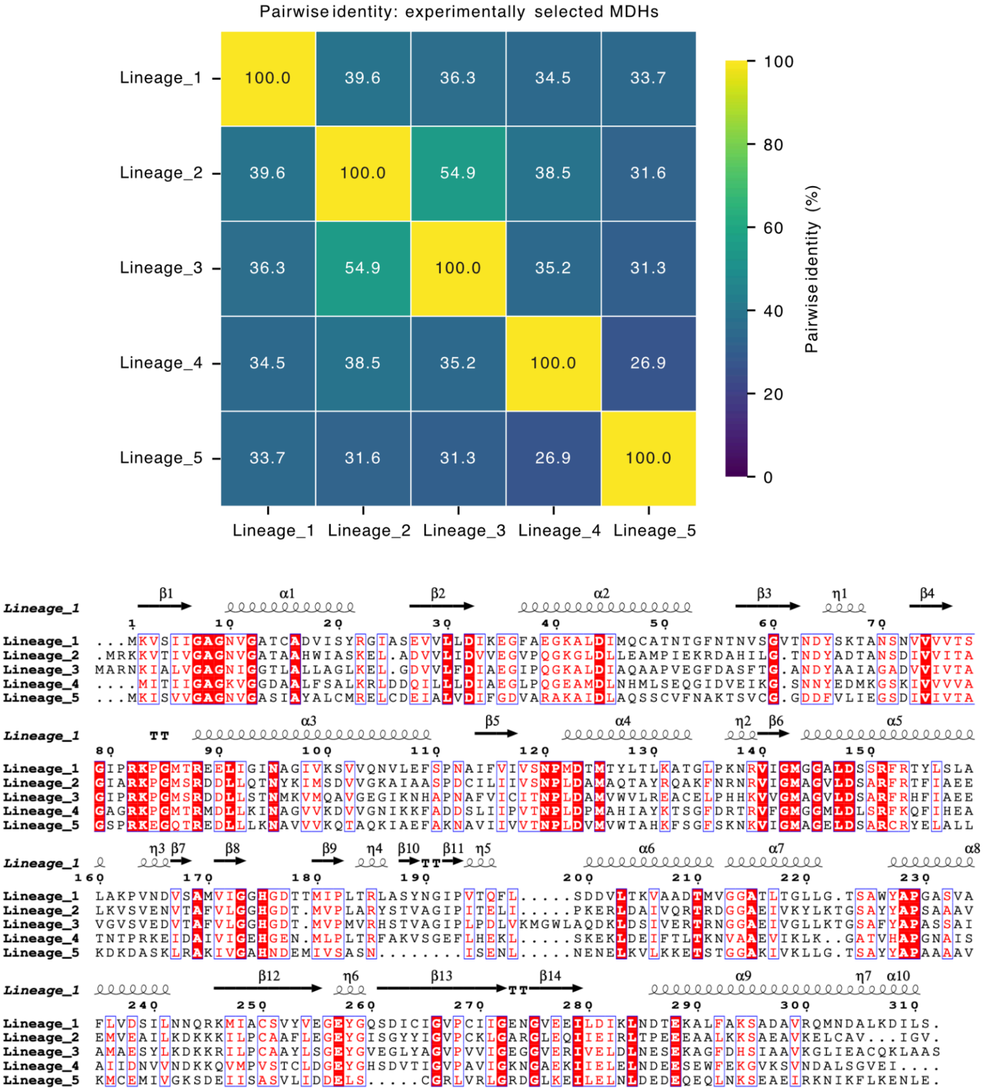


**Supplementary Figure 16.** Sequence comparison of experimentally selected malate dehydrogenases. Multiple-sequence alignment of the five selected malate dehydrogenase (MDH) sequences generated using Clustal Omega[7] and visualised with ESPript[8]. The pairwise sequence-identity matrix was calculated from the uncorrected Clustal Omega distances as 100 * (1-d). Values represent per cent amino-acid identity, with colours indicating identity according to the viridis scale. The mean identity across all ten unique sequence pairs was 36.26%. Secondary-structure elements are displayed above the alignment and were derived from structure predicted by AlphaFold2.

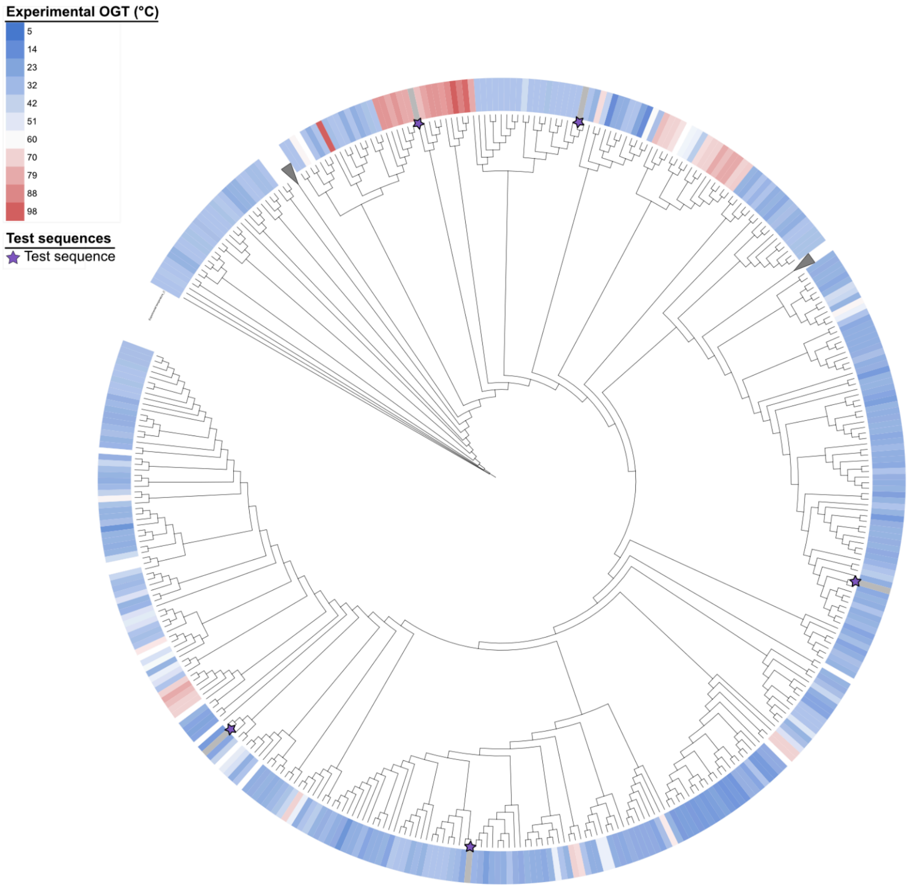


**Supplementary Figure 17.** Phylogenetic distribution of experimentally characterised lineages within the MDH family that form the test set. Phylogenetic tree of malate dehydrogenase (MDH) family sequences. The outer heat map indicates the experimentally determined optimal growth temperature (OGT) associated with each sequence, with colours ranging from blue for lower temperatures to red for higher temperatures. Purple stars indicate the sequences selected as experimental lineages. Two large clades were collapsed for visualisation and contain 741 and 576 sequences, respectively.

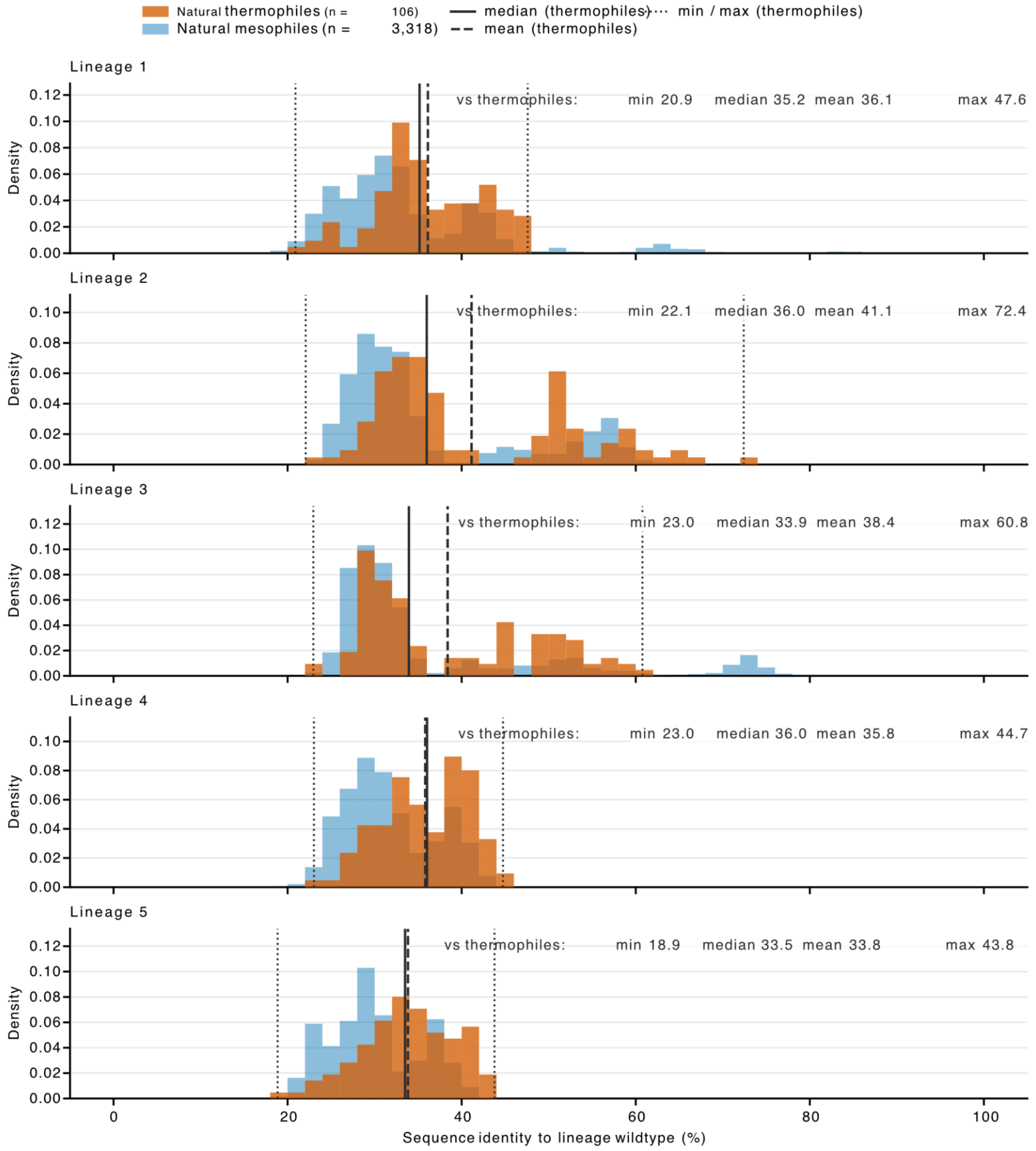


**Supplementary Figure 18.** The five MDH lineage wildtypes are only remotely related to natural thermophilic homologues. Distribution of pairwise amino-acid sequence identity between each lineage wildtype (Lineage 1–5) and natural COG0039 malate dehydrogenases of known environmental temperature. Thermophiles (orange, n = 106) are sequences whose source organism has a measured environmental temperature ≥ 60 °C (median 70 °C, maximum 98 °C); mesophiles (blue, n = 3,318, OGT 4–59 °C) are shown as a reference distribution. Identity was computed from independent global pairwise alignments (BLOSUM62, gap open −11, gap extend −1, terminal gaps unpenalised) as the number of identical residues divided by the length of the shorter ungapped sequence. Both distributions are plotted as densities, as the two sets differ ~30-fold in size. Vertical lines mark the minimum and maximum (dotted), median (solid) and mean (dashed) of the thermophile distribution; the same values are printed in each panel.

#### ZeroShot THOR activity progress curves per variant (30 °C)

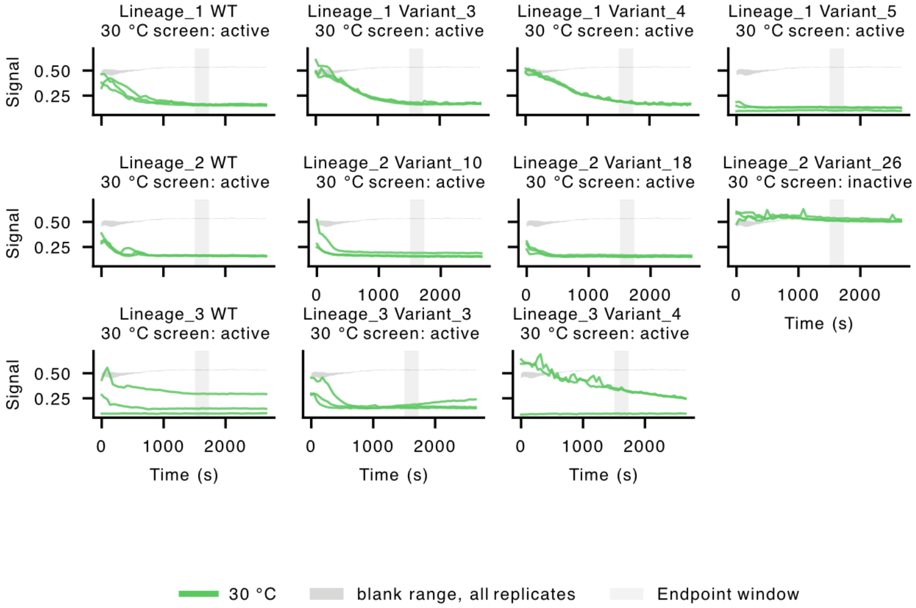


**Supplementary Figure 19.** Activity progress curves for wild-type enzymes and THOR ZeroShot-designed variants at 30 °C. Each panel shows an individual enzyme preparation, with lineage and variant identifiers indicated above. Lines show raw, unnormalised assay signals from three replicate wells. Primary and backup preparations are displayed separately where available. The grey envelope indicates the minimum–maximum range of blank-control readings, and the pale vertical band marks the endpoint window used for activity screening (1530–1710 s). Panel annotations report activity classifications based on the 30 °C screen, using the endpoint and rate criteria described in Methods.

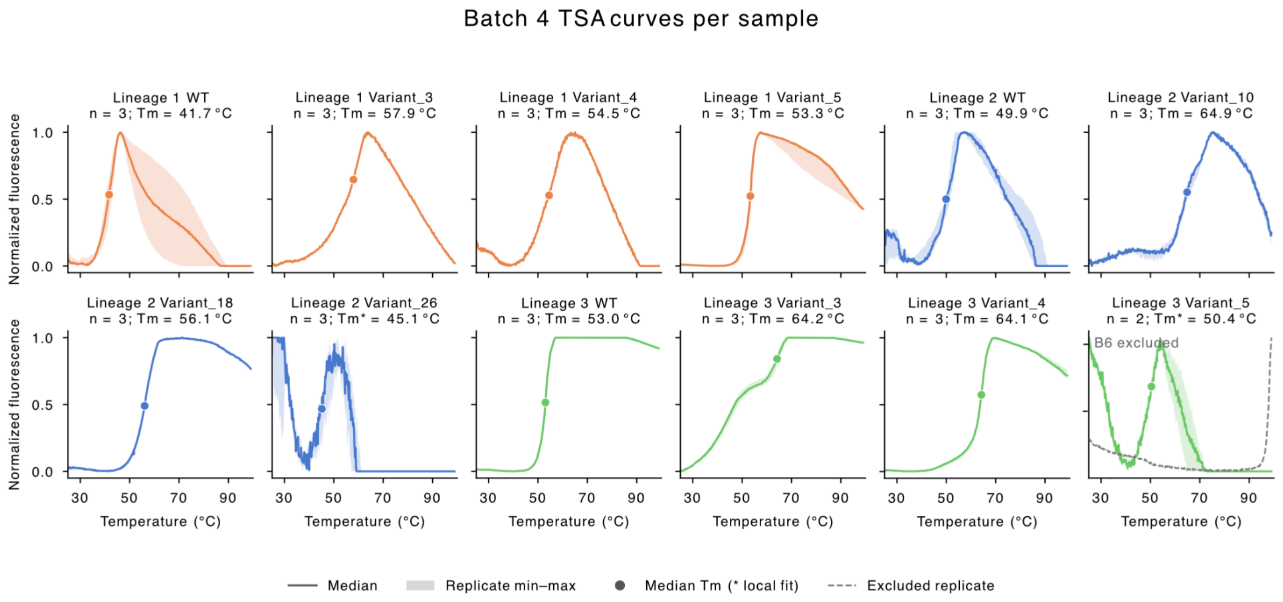


**Supplementary Figure 20.** Thermal shift assay profiles of wildtypes and THOR ZeroShot variants. Normalised fluorescence profiles are shown for three wildtypes (WT) and eight variants from Lineages 1–3. Fluorescence was normalised separately for each well using its minimum signal preceding or at the fluorescence peak and its peak signal, with values restricted to 0–1. Solid lines show the pointwise median, and shading indicates the minimum–maximum range across wells. Where temperature grids differed, curves were interpolated within their shared measured temperature interval before aggregation. Samples were measured in three replicate wells, except for the Lineage 1 WT (six wells). Circles and panel titles indicate median per-well fitted melting temperatures (*T*□). Orange, blue, and green denote Lineages 1, 2, and 3, respectively.

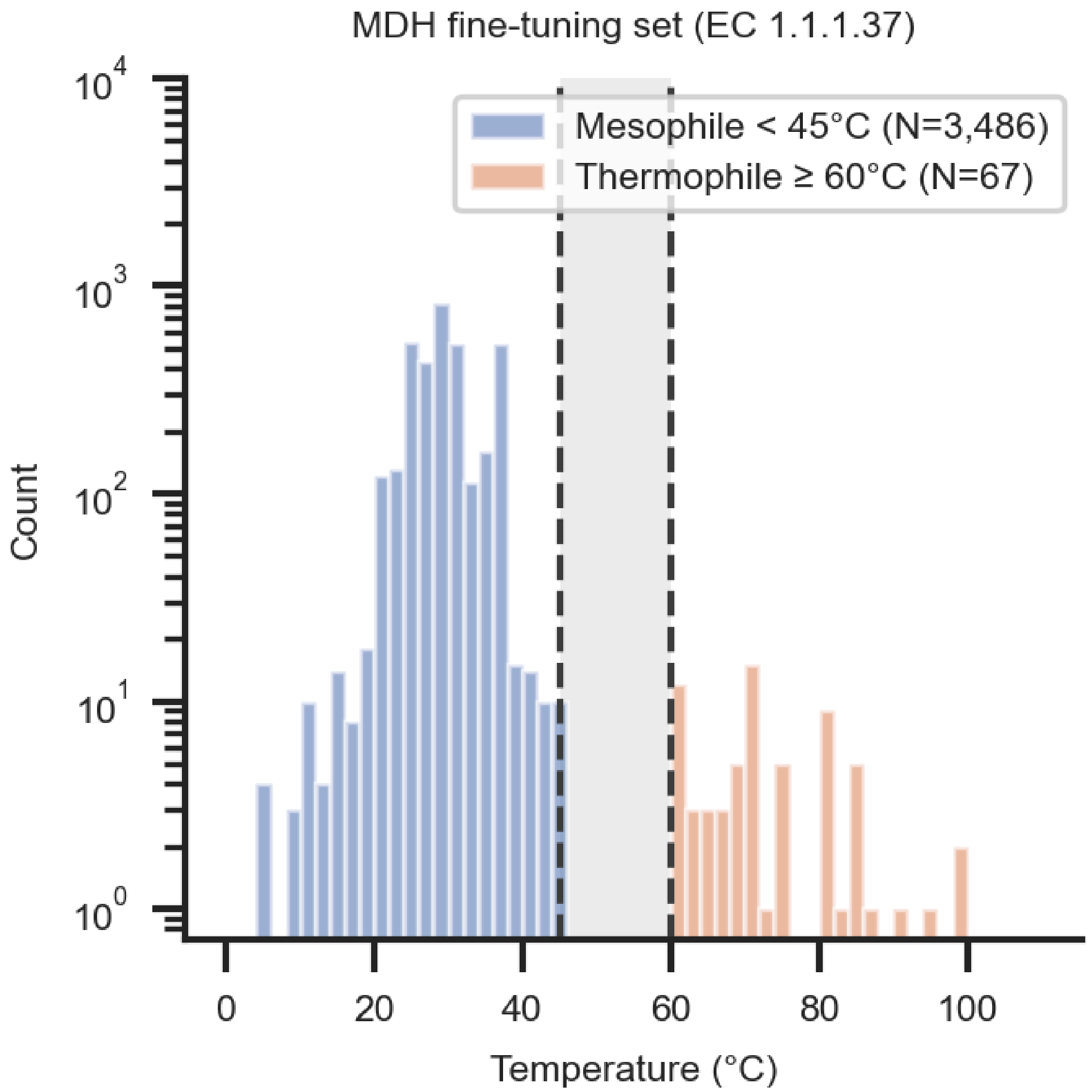


**Supplementary Figure 21**. Temperature distribution of the MDH fine-tuning set. Malate dehydrogenase sequences (EC 1.1.1.37) drawn from the OGT dataset, in which every sequence carries a measured source-organism growth temperature. Sequences were split into mesophiles below 45 °C (blue, N = 3,486) and thermophiles at or above 60 °C (orange, N = 67); the intervening 45–60 °C band (shaded) was excluded so that the two classes are separated by a 15 °C margin, removing a further 75 sequences. Bins are 2 °C wide and the count axis is logarithmic. Median growth temperature is 28 °C for the mesophiles and 70 °C for the thermophiles. All 3,628 sequences matching the EC filter also carry COG0039 at root level, so the functional selection is unambiguous.

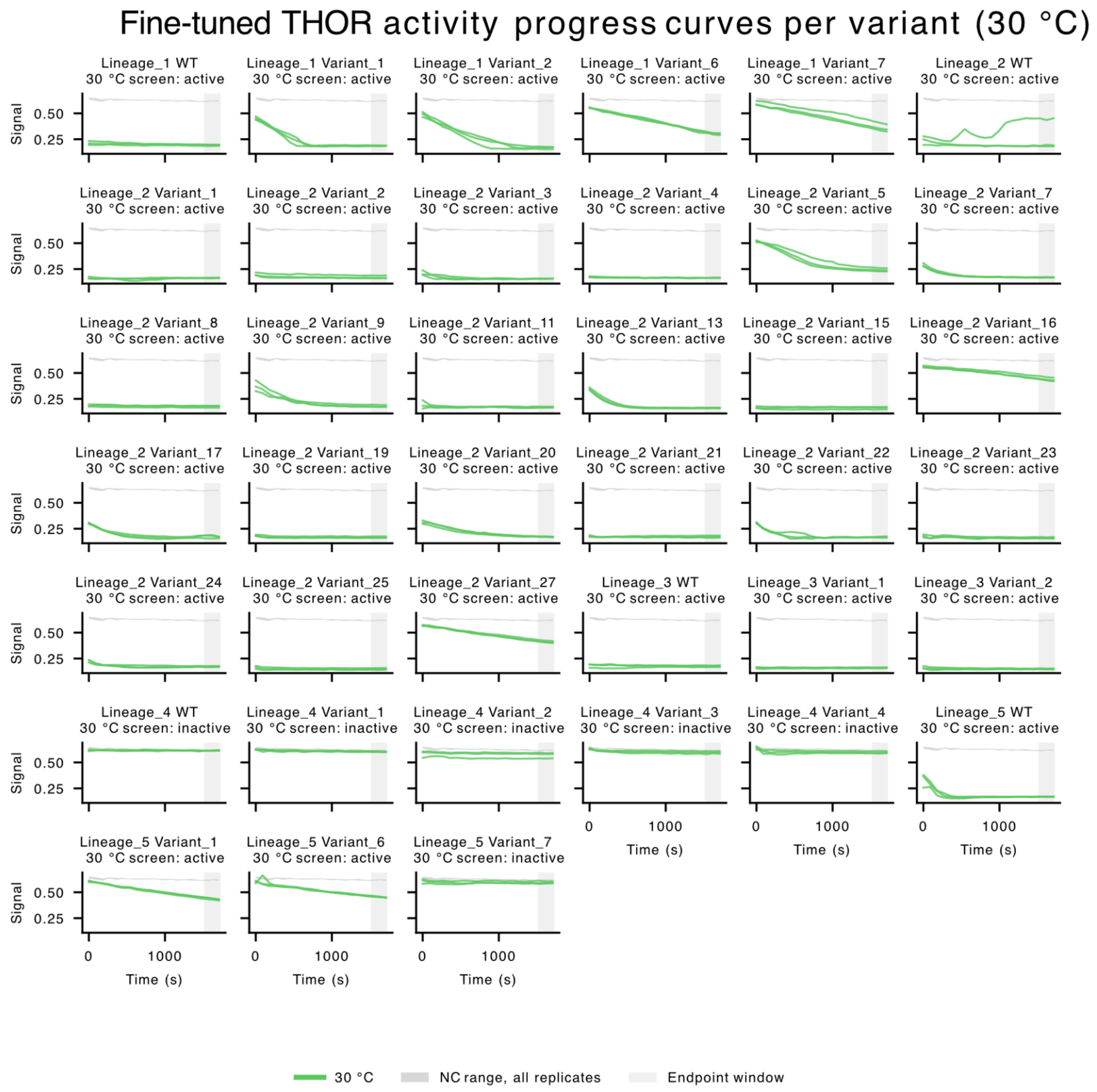


**Supplementary Figure 22.** Activity progress curves for wild-type enzymes and THOR fine-tuned designed variants at 30 °C. Each panel shows an individual enzyme preparation, with lineage and variant identifiers indicated above. Lines show raw, unnormalised assay signals from three replicate wells. Primary and backup preparations are displayed separately where available. The grey envelope indicates the minimum–maximum range of blank-control readings, and the pale vertical band marks the endpoint window used for activity screening (1530–1710 s). Panel annotations report activity classifications based on the 30 °C screen, using the endpoint and rate criteria described in Methods.

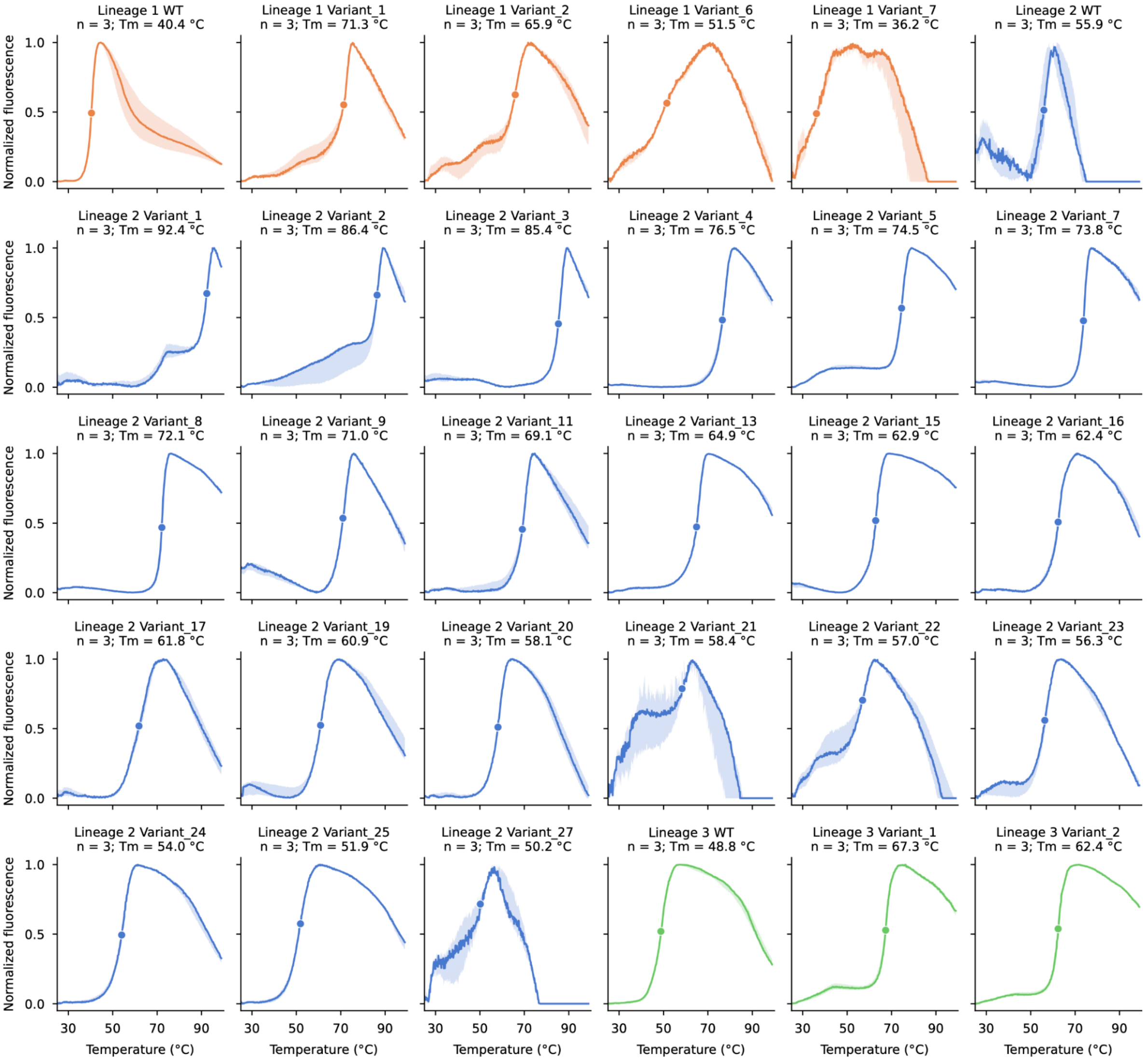


**Supplementary Figure 23.** Thermal shift assay profiles of wildtypes and Fine-Tuned THOR variants. Normalized fluorescence profiles are shown for three wildtypes (WT) and 27 variants from Lineages 1–3. Each panel represents one sample measured in three replicate wells. Fluorescence was normalized separately for each well by subtracting the minimum signal preceding or at the fluorescence peak and dividing by the peak-to-minimum difference; values were restricted to 0–1. Solid lines show the pointwise median, and shaded regions indicate the minimum–maximum range across wells. Circles mark the median of the per-well fitted melting temperatures (*T*□), also reported in the panel titles. Orange, blue, and green denote Lineages 1, 2, and 3, respectively. Variant numbering is specific to each lineage.

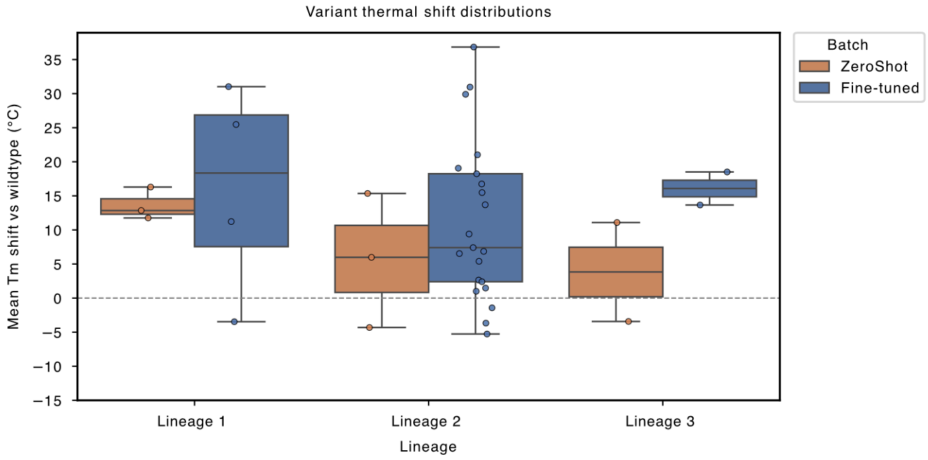


**Supplementary Figure 24.** Experimental thermal stability shifts of generated MDH variants. Distribution of experimentally measured melting temperature shifts (ΔTm) for generated variants from three MDH lineages, determined using thermal shift assays (TSA). ΔTm is reported relative to the corresponding wild-type sequence, with positive values indicating increased thermal stability. Variants were generated using either the **ZeroShot** model, which was trained without orthologous sequences from the corresponding MDH family, or the **fine-tuned** model, which was additionally trained on a small set of orthologous sequences. The dashed horizontal line indicates no change relative to the wild type (ΔTm = 0 °C).

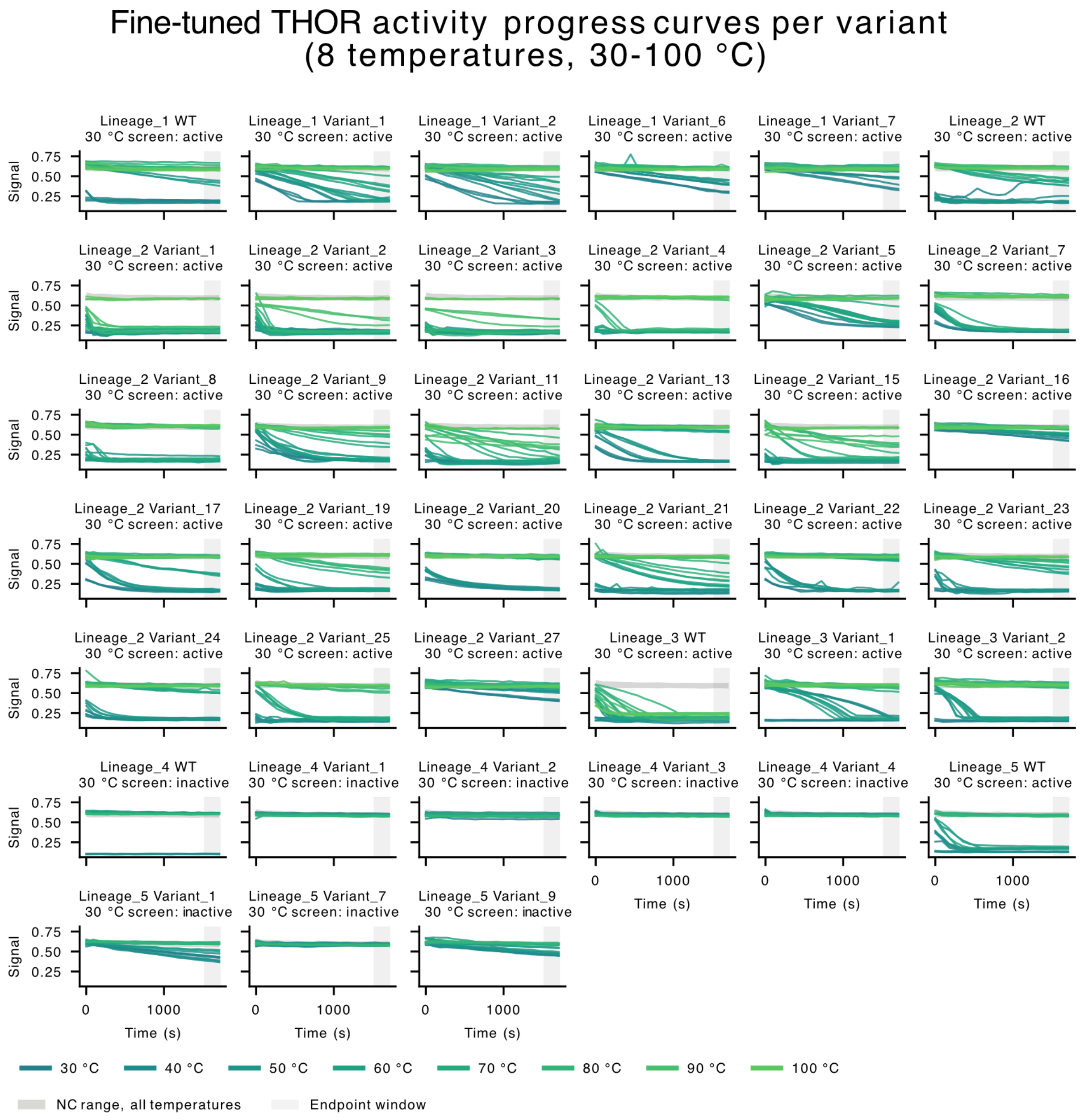


**Supplementary Figure 25.** Activity progress curves for wild-type enzymes and THOR fine-tuned designed variants across assay temperatures. Each panel shows an individual enzyme, with lineage and variant identifiers indicated above. Raw, unnormalised assay signals are plotted against time for three replicate wells per temperature; colours indicate temperatures from 30 to 100 °C in 10 °C increments. Lineage 3 and Lineage 4 enzymes were measured only at 30–80 °C. The grey envelope shows the minimum–maximum range of negative-control readings pooled across temperatures. The pale vertical band marks the endpoint window used for activity screening (1530–1710 s). Panel annotations report activity classifications based exclusively on the 30 °C screen, using the endpoint and rate criteria described in Methods.

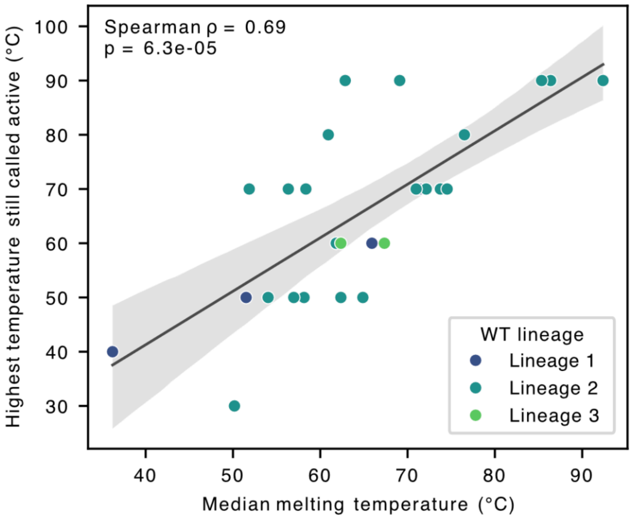


**Supplementary Figure 26.** Relationship between melting temperature and retention of enzymatic activity. For each of the 25 variants represented in both assays, the median melting temperature measured by TSA was compared with the highest assayed treatment temperature at which the variant retained at activity. Points are colored according to the parental wild-type lineage, and the line shows the overall linear regression. A modest positive correlation was observed (Spearman's = 0.69, p = 6.3e-5). Variants retaining activity at the highest assayed temperature are right-censored because their maximum functional temperature may exceed the experimental range.

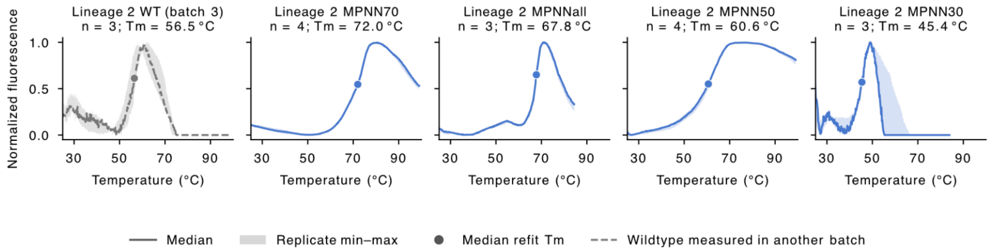


**Supplementary Figure 27. Thermal shift assay profiles of Batch 5 ProteinMPNN designs.** Normalized fluorescence profiles are shown for the four Lineage 2 designs MPNN30, MPNN50, MPNN70, and MPNNall. Fluorescence was normalized separately for each well using its minimum signal preceding or at the fluorescence peak and its peak signal, with values restricted to 0–1. Solid lines show the pointwise median, and shaded regions indicate the minimum–maximum range across wells after interpolation within their shared measured temperature interval. MPNN30 and MPNNall were measured in three replicate wells each; MPNN50 and MPNN70 were measured in four wells each.

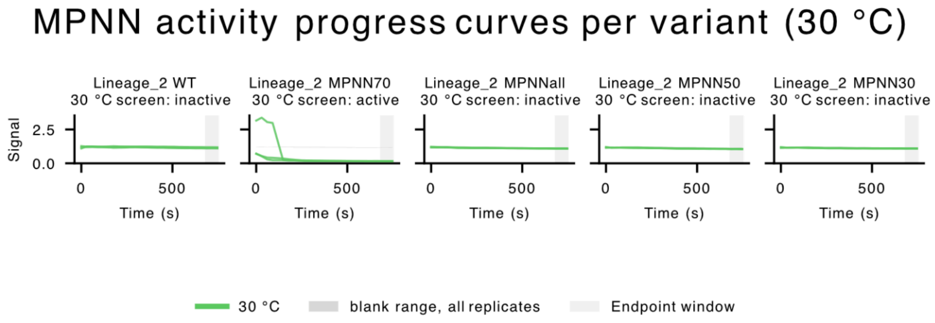


**Supplementary Figure 28.** Activity progress curves for wild-type Lineage 2 and MPNN designed variants at 30 °C. Each panel shows an individual enzyme preparation, with lineage and variant identifiers indicated above. Lines show raw, unnormalised assay signals from three replicate wells. Primary and backup preparations are displayed separately where available. The grey envelope indicates the minimum–maximum range of blank-control readings, and the pale vertical band marks the endpoint window used for activity screening (687–747 s). Panel annotations report activity classifications based on the 30 °C screen, using the endpoint and rate criteria described in Methods.

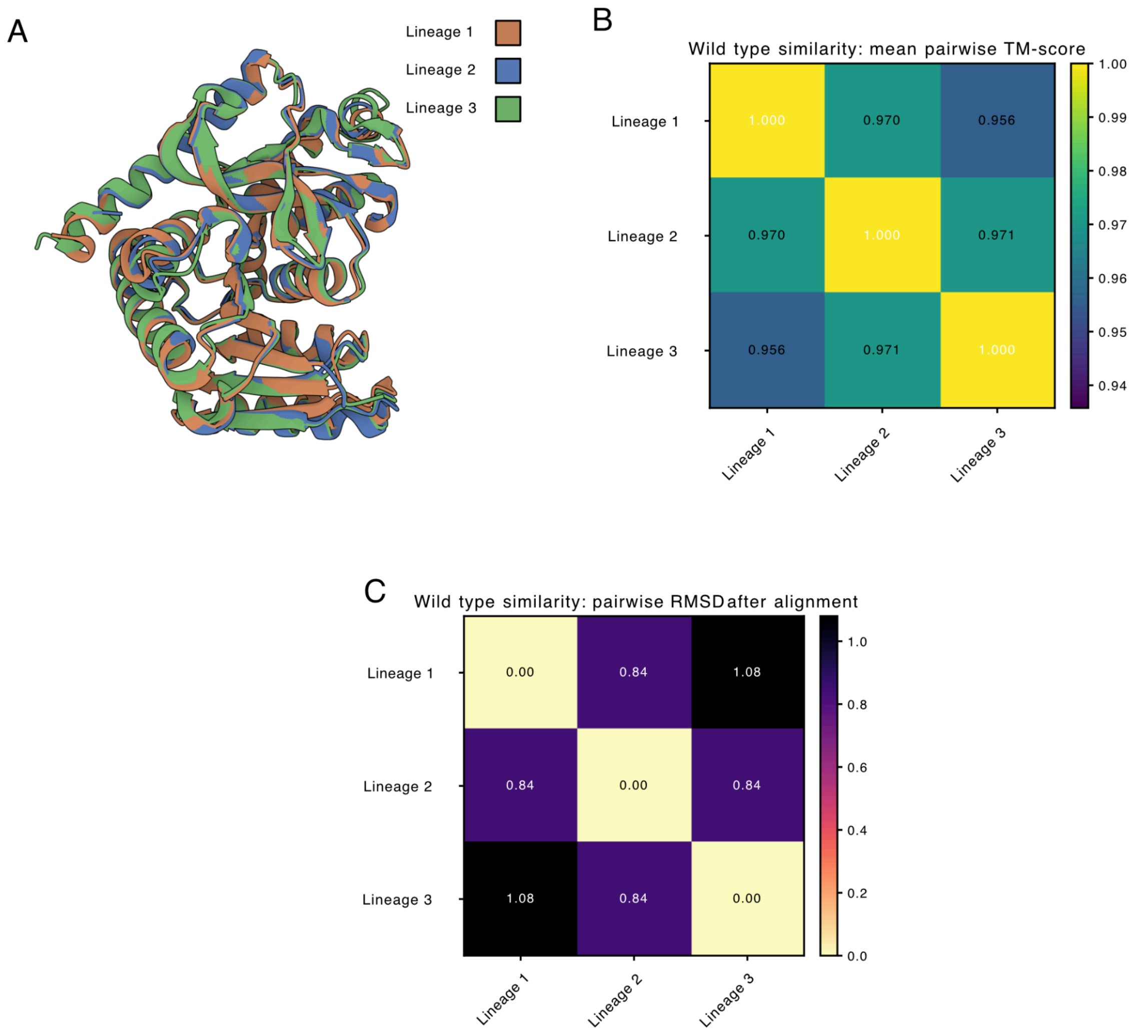


**Supplementary Figure 29.** Structural comparison of wild-type proteins from Lineages 1–3.
(A) Structural superposition of the wild-type protein structures from Lineage 1 (orange), Lineage 2 (blue), and Lineage 3 (green) following alignment with TM-align. (B) Pairwise structural similarity between the three wild-type structures, reported as TM-scores. (C) Pairwise root-mean-square deviation (RMSD; Å) between the structures following alignment. The high TM-scores and low RMSD values indicate that the wild-type proteins from all three lineages adopt highly similar overall structures.

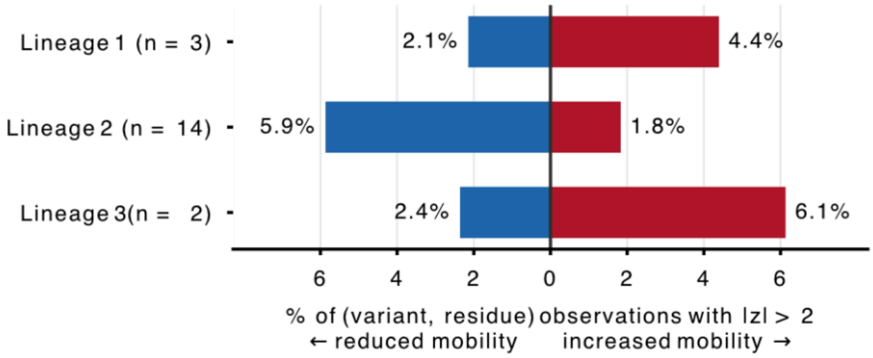


**Supplementary Figure 30.** Direction of RMSF change relative to wild type, by lineage.
For each simulated system, per-residue RMSF was computed from four independent MD replicas (three covering 0–300 ns and one covering 0–500 ns, frames stored every 20 ps) and averaged across replicas. For every variant, ΔRMSF was taken as the replica-averaged variant RMSF minus that of its own lineage wild type, and the resulting ΔRMSF profile was converted to Z-scores across residues within each variant. Bars show the percentage of (variant, residue) observations in each lineage with $Z > 2$ (red, right: mobility increased relative to wild type) and $Z < -2$ (blue, left: mobility reduced). Lineage 1, n = 3 variants over 311 residues; lineage 2, n = 14 over 308; lineage 3, n = 2 over 318. Because the denominator is the number of variants multiplied by the number of residues, the percentages are mean per-variant rates and are therefore comparable between lineages despite the differing numbers of variants per lineage. Lineages 1 and 3 are biased towards increased mobility (4.4% and 6.1% of observations with $Z > 2$, versus 2.1% and 2.4% with $Z < -2$), whereas lineage 2 is dominated by reduced mobility (5.9% with $Z < -2$ versus 1.8% with $Z > 2$). Z-scores are descriptive within-profile scores, not significance tests, and no multiple-testing correction is applied. Only variants with an experimentally measured ΔTm of at least +5 °C relative to their wild type were simulated.

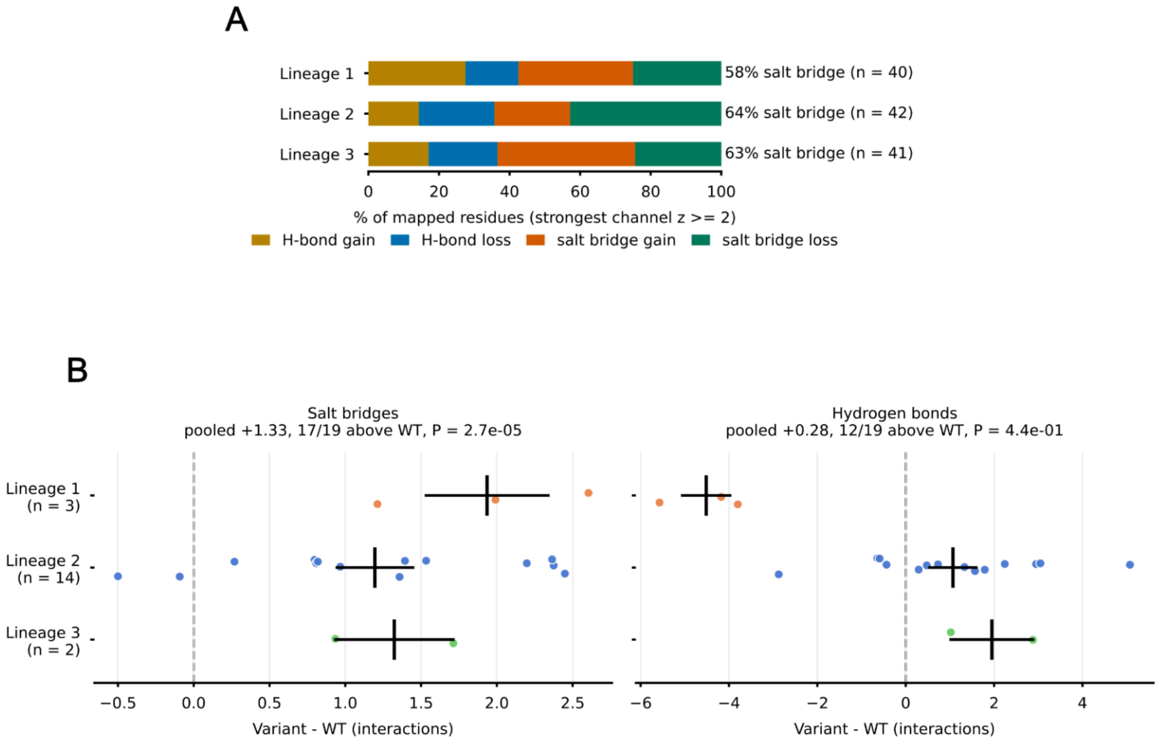


**Supplementary Figure 31**. Interaction-class composition and absolute interaction changes relative to wild type.(A) Interaction class underlying the mapped structural signal. For each wild-type residue, four wild-type-relative interaction channels were formed: hydrogen-bond gain, hydrogen-bond loss, salt-bridge gain and salt-bridge loss. each averaged over all variants in the lineage and then converted to Z-scores across residues within that lineage. A residue was counted as mapped when its strongest channel reached $Z \geq 2$, and was attributed to that channel; this is the construction used to color Fig. 4B. Bars give the channel composition of the mapped residues in each lineage (gold, hydrogen-bond gain; blue, hydrogen-bond loss; orange, salt-bridge gain; teal, salt-bridge loss), with the salt-bridge share and the number of mapped residues given at right (40, 42 and 41 mapped residues of 311, 308 and 318 respectively). All three lineages leaned similarly towards salt-bridge-associated regions (57.5%, 64.3% and 63.4%), with no evidence of a difference between them ($\chi^2 = 0.47$, df = 2, P = 0.79). Because the Z-scores are computed within each profile, this panel reports where interaction signal concentrates and is insensitive to its magnitude. (B) Absolute interaction counts relative to wild type. Each point is one variant, plotted as the mean of its four replica means minus that of its own wild type; the vertical bar marks the lineage mean and the horizontal bar its standard error, and the dashed line marks no change from wild type. Salt bridges increased in all three backgrounds (17 of 19 variants above wild type; pooled mean +1.33, Wilcoxon P = $2.7 \times 10^{-5}$), with no difference in the size of the gain between lineages (Kruskal-Wallis P = 0.40). Hydrogen bonds showed no overall shift (pooled mean +0.28, P = 0.44) but differed between backgrounds (Kruskal-Wallis P = 0.024): lineage 1 variants lost hydrogen bonds (−4.5, one-sample t-test P = 0.014) whereas lineages 2 and 3 gained them (+1.1 and +2.0). Tests treat the four replica means as the independent units. Z-scores in (A) are descriptive within-profile scores rather than significance tests, and no multiple-testing correction was applied.

### Supplementary tables

**Supplementary Table 1.** Recall and precision characteristics of Environmental Temperature (ET) regression model for temperature cutoffs at 45, 60, and 70 degrees.

| temperature | >=45 C | >=60 C | >60 C | >=70 C |
|---|---|---|---|---|
| recall | 0.71507761 | 0.71932638 | 0.82217573 | 0.57469244 |
| precision | 0.45679887 | 0.69805447 | 0.61167315 | 0.8034398 |
| F-score | 0.55747623 | 0.70853081 | 0.70147256 | 0.55747623 |

**Supplementary Table 2**. Composition of the ten enzyme test families. The ten EggNOG root orthologous groups (COGs) making up the held-out enzyme test set used in Fig. 2. Sequences were assigned to families by eggNOG-mapper; the EC column gives the most frequent EC annotation within each family. The test set combines two sources: sequences from IMG thermophile genomes, all of which derive from source organisms with an optimal growth temperature (OGT) ≥ 60 °C, and UniProt enzymes carrying the mesophilic class label. The thermophile column reports the IMG fraction of each family. Lengths are median [interquartile range] in residues; all test sequences fall within the 512-residue limit of the temperature predictor, so no family is affected by length-based exclusion. Families differ substantially in thermophile representation, from 53.9% (COG0673) down to 0.6% (COG4188), which is the family used as the no-natural-thermophile case in the phylogenetic enrichment analysis (Supplementary Fig. SX). Across all ten families the test set comprises 450,252 sequences, 110,215 (24.5%) of them thermophilic.

| **root_COG** | **Family** | **EC** | **PFAM** | **n_sequences** | **n_thermophile** |
|---|---|---|---|---|---|
| COG1012 @1\|root | Acyl-CoA reductase | 1.2.1.18,1.2.1.27 | Aldedh | 107796 | 27764 |
| COG1028 @1\|root | FabG/SDR dehydrogenase | 1.1.1.100 | adh_short_C2 | 140862 | 26397 |
| COG0108 @1\|root | RibB / DHBP | 3.5.4.25,4.1.99.12 | DHBP_synthase,GTP_cyclohydro2 | 29812 | 7234 |
| COG0062 @1\|root | Nnr/NAXE | 4.2.1.136,5.1.99.6 | Carb_kinase,YjeF_N | 20527 | 7582 |
| COG0065 @1\|root | LeuC | 4.2.1.33,4.2.1.35 | Aconitase | 35679 | 11863 |
| COG0039 @1\|root | MDH | 1.1.1.37 | Ldh_1_C,Ldh_1_N | 52942 | 7646 |
| COG0673 @1\|root | MviM-like dehydrogenase | 1.1.1.18,1.1.1.369 | GFO_IDH_MocA,GFO_IDH_MocA_C | 21428 | 11543 |
| COG3842 @1\|root | ABC transporter ATPase | 3.6.3.31 | ABC_tran,TOBE_2 | 24967 | 9793 |
| COG2032 @1\|root | SODC | 1.15.1.1 | Sod_Cu | 13045 | 374 |
| COG4188 @1\|root | α/β-hydrolase | 3.1.1.47 | PAF-AH_p_II | 3194 | 19 |

A

kinetics parameters, 30 °C

| Variant | Km (mM) | kcat (s^-1) | kcat/Km (mM^-1 s^-1) |
|---|---|---|---|
| WT | 0.0361 | 11.451 | 316.953 |
| Variant 2 | 0.0573 | 31.051 | 542.138 |
| Variant 1 | 0.1565 | 16.205 | 103.562 |
| MPNN70 | 0.0112 | 0.969 | 86.833 |

B

kinetics parameters, 70 °C

| Variant | Km (mM) | kcat (s^-1) | kcat/Km (mM^-1 s^-1) |
|---|---|---|---|
| WT | 0.0766 | 64.640 | 843.548 |
| Variant 2 | 0.1864 | 85.596 | 459.167 |
| Variant 1 | 0.1669 | 38.622 | 231.457 |
| MPNN70 | 0.0151 | 5.096 | 338.376 |

C

kinetics parameters, 90 °C

| Variant | Km (mM) | kcat (s^-1) | kcat/Km (mM^-1 s^-1) |
|---|---|---|---|
| WT | 0.0646 | 10.102 | 156.293 |
| Variant 2 | 0.0747 | 26.988 | 361.124 |
| Variant 1 | 0.5192 | 19.335 | 37.240 |
| MPNN70 | 0.0305 | 1.757 | 57.679 |

**Supplementary Table 3.** Steady-state kinetic parameters for Lineage 2 malate dehydrogenase variants. Michaelis–Menten parameters for the Lineage 2 wild type (WT), the THOR designs Variant 1 and Variant 2, and the ProteinMPNN control MPNN70 at (A) 30 °C, (B) 70 °C and (C) 90 °C. Initial rates were measured from NADH absorbance at 340 nm across 6–14 substrate concentrations (0.01–2.0 mM, three to five replicates each) and fitted by non-linear least squares. $k_{cat}$ is normalised to enzyme concentration and so is comparable across variants despite the higher loading used for MPNN70. Intervals are wide where the substrate range did not approach saturation, most notably for Variant 1 at 70 and 90 °C, where $k_{cat}/K_m$ is better determined than either parameter alone.
